# Insights into the role of Nup62 and Nup93 in assembling cytoplasmic ring and central transport channel of the nuclear pore complex

**DOI:** 10.1101/2022.02.28.482420

**Authors:** Pankaj K. Madheshiya, Ekta Shukla, Jyotsna Singh, Shrankhla Bawaria, Mohammed Yousuf Ansari, Radha Chauhan

## Abstract

The nuclear pore complex (NPC) is a highly modular assembly of 34 distinct nucleoporins (Nups), to form a versatile transport channel between the nucleus and cytoplasm. Among them, Nup62 is known as an essential component for nuclear transport while, Nup93 for the proper nuclear envelope assembly. These Nups constitute various NPC subcomplexes: such as central transport channel (CTC), cytoplasmic ring (CR) and inner ring (IR). However, how they play their role in the NPC assembly and transport activity is not clear. Here we delineated the interacting regions, conducted biochemical reconstitution and structural characterization of the mammalian CR complex to reveal its intrinsic dynamic behaviour and a distinct ‘4’ shaped architecture resembling the CTC complex. Our data demonstrate that Nup62 coiled-coil domain is critical to form both Nup62•Nup88 and Nup62•Nup88•Nup214 heterotrimers and both can bind to the Nup93. We therefore propose that Nup93 act as a ‘sensor’ to bind to Nup62 shared heterotrimers including Nup62•Nup54 heterotrimer of the CTC, which was not shown previously as an interacting partner. Altogether, our study establishes that the Nup62 via its coiled-coil domain is central to form compositionally distinct yet structurally similar heterotrimers, and the Nup93 anchors these diverse heterotrimers by recognizing them non-selectively.

## INTRODUCTION

Nuclear pore complexes (NPCs) function as the exclusive gateways between the nucleus and the cytoplasm to facilitate the bi-directional nucleocytoplasmic transport (Beck and Hurt 2017; Rout et al. 2000). The NPCs are highly modular and intricate structures ranging from ~60 MDa in yeast to ~120 MDa in humans (Alber, Dokudovskaya, Veenhoff et al. 2007; Cronshaw et al. 2002) and composed of about 34 distinct nucleoporins (Nups), which are present in multiple copies to form an eight-fold rotational symmetric core across the nucleo-cytoplasmic axis. These Nups are arranged in various sub-complexes to carry out the distinct biologically conserved function such as mRNPs export into the cytoplasm and transport of cargoes into and out of the nucleus (Hoelz et al. 2011; Grossman et al. 2012; Lin and Hoelz 2019; Schwartz 2017; Beck and Hurt 2017). Although radially symmetric, NPC shows nuclear-cytoplasmic asymmetry and is composed of three structural features: a nuclear ring (NR) with a nuclear basket that extends into the nucleoplasm, a central transport channel (CTC) along with inner ring Nups and a cytoplasmic ring (CR) with filaments which stretch out into the cytoplasm (Fig. S1). The detailed knowledge of NPC structure is a prerequisite for mechanistic understanding of its function. The unusually large size of the NPC, together with its conformational plasticity, represents a challenge for its 3D structure determination at atomic resolution. Moreover, the complete interaction network of NPC components and their biochemical behaviour is still not fully understood. These bottlenecks have significantly hampered the basis for modular NPC assembly and its role in versatile transport functions.

Throughout the vertebrates, a major component of the cytoplasmic ring (CR) is the Nup88 complex harbouring three proteins Nup62, Nup88, and Nup214 (Fig. S1). Both Nup88 and Nup214 possess an N-terminal β-propeller domain followed by an α-helical region (Fig. 1a and 1c). Additionally, Nup214 contains an extensive unstructured FG repeat region at its C-terminal (Fig. 1c). On the other hand, Nup62 possesses an N-terminal unstructured region with FG repeats and C-terminal α-helical region (Fig. 1b). These three proteins in Nup88 complex (Nup88•Nup62•Nup214) are suggested to interact via coiled-coil domains (Huang, Zhang, Zhu, Zeng et al. 2020). Various mRNA export factors that are important for messenger ribonucleoprotein particles (mRNPs) remodelling; bind at the CR of the NPC prior to translation (Bailer et al. 2000; Hutten and Kehlenbach 2006; Kalverda, Pickersgill et al. 2010; Napetschnig 2007; Snay-hodge et al. 1998). Moreover, Nup214 and Nup88 are also associated with various human neurological disorders, cardiac diseases and cancers (Lin and Hoelz 2019; Yarbrough et al. 2014; Hurt and Alwin 2010; Nofrini et al. 2016; Ciomperlik et al. 2016). However, the detailed mechanistic function of these Nups with the diseases is not characterised. The interaction studies of the Nup88 complex with other neighbouring Nups are limited, except for reports from lower eukaryotes like *S. cerevisiae* and *C. thermophilum*. The homolog of the Nup88 complex in fungi is known as the Nup82 complex containing Nup82 (Nup88), Nup159 (Nup214) and Nsp1 (Nup62). The cross-linking mass spectrometry data suggests the inter-domain interaction network between the subunits of Nup82 complex and with the linker Nups (such as Nup98, Nup145C and Nup145N) that are responsible for the NPC assembly (Gaik, Dirk, Von Appen et al. 2015; Teimer et al. 2017). Moreover, X-ray crystallographic structures of the fragments of some Nups and their complexes are available, such as Nup82^NTD^•Nup159^Tail^ with Nup116^CTD^, Nup145N and Nup98^APD^ (Stuwe et al 2012; Stuwe, Correia et al. 2016; Napetschnig 2007; Stuwe, Bley, Thierbach, Petrovic et al. 2016; Yoshida et al. 2011). The negative stain electron microscopy (EM) of the fungal Nup82 complex has revealed an overall ‘P’ shape or ‘D’ shape architecture for this complex (Gaik, Dirk, Appen et al. 2015; Teimer et al. 2017). However, in case of metazoans, low resolution Cryo-electron tomographic (Cryo-ET) reconstructions of the *X. laevis* CR (Huang, Zhang, Zhu, Zeng et al. 2020) showing the positioning of Nup88•Nup62•Nup214 complex along with the IR complex (Nup93, Nup205, Nup188, Nup155 and Nup35) and Y-shaped complex (Nup107, Nup133, Sec13, Seh1, Nup160, Nup43, Nup96, Nup75 and Nup37) is available. Similar tomography maps from *H. sapiens* (Bui, Appen et al. 2013; Hurt and Beck 2015; Appen, Kosinski, Sparks et al. 2015; Kosinski, Mosalaganti, Appen, et al. 2016; Schuller, Wojtynek et al. 2021) are available lately showing proximity of CR, IR and Y shape complexes. However, in these cases, precise inter-domain interactions and biochemical behaviour of CR and IR complexes have not been understood yet. The limitations in purifying these stable complexes have impeded the attempts to pursue structural studies for the mammalian complexes.

**Figure 1:**
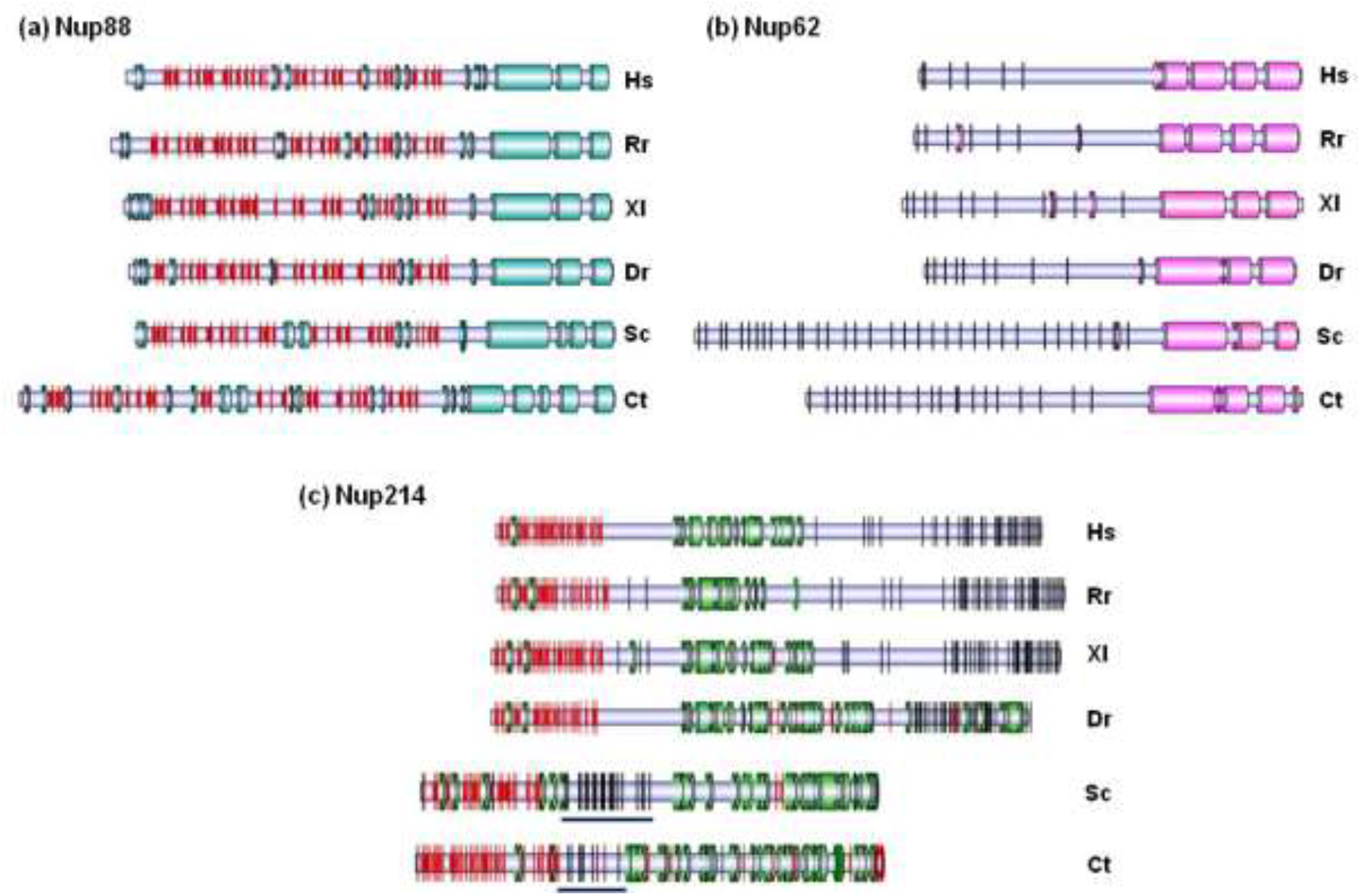
Secondary structure organization of the Nup88, Nup62 and Nup214 sequences. Organization of secondary structure domains for Nup88 (a), Nup62 (b) and Nup214 (c) from species: *Hs- Homo sapiens, Rn- Rattus norvegicus, Xl- Xenopus laevis, Dr- Danio rerio, Sc- Saccharomyces cerevisiae, Ct- Chaetomium thermophilum*. Predicted α-helical domains are represented by cylinder and β-sheets by red arrow. FG-repeats in the sequences are represented by vertical black lines. The FG repeats sandwiched between the helical region and β-propeller region in Nup214 sequences of *Saccharomyces* and *Chaetomium* are underlined in black.

Recent studies have demonstrated the diverse nature of Nups in different species (Chopra et al. 2019; Eibauer et al. 2015; Kim, Fernandez-Martinez, Nudelman, Shi, Zhang et al. 2018; Mosalaganti et al. 2018; Appen, Kosinski, Sparks et al. 2015; Field and Rout, 2019). There are significant differences in the arrangement and architecture of CR Nups in different organisms. Evidence from the vertebrate *X. laevis*, shows that the Nup88 complex adopts a ‘rake-shaped’ structure which is different from fungi where corresponding homolog’s two copies (Nup82 complex) are arranged in a parallel fashion (Tai, Zhu, Ren et al. 2022). In another study (unpublished data, Bley, Liu, Nie, Mobbs, Petrovic, Gres et al. 2021), it is shown that both yeast and corresponding human heterotrimeric Nup88 complexes have a “P” shaped structure. Further, the mechanisms for the attachment of the CR complexes in *C. thermophilum* NPC is distinct from the vertebrates where Nup358 is wrapped around the CR to form cytoplasmic filaments (unpublished data, Tai, Zhu, Ren et al. 2021).

As Nup88•Nup62•Nup214 complex is central to the CR, we explored the detailed interaction studies followed by biochemical reconstitution of this mammalian complex to understand the role of individual Nups. To characterise the interaction networks, we performed the *in-silico* protein-protein interactions prediction using CoRNeA platform (Chopra et al. 2020) coupled with *in-vitro* tandem affinity pull down (TAP) assays and dissected the inter-domain interactions among these three Nups. Our study revealed that the α-helical coiled-coil regions of these proteins are critical to form stable complexes of Nup88•Nup62 and Nup88•Nup62•Nup214. Further, we purified these complexes and performed size exclusion chromatography coupled with multiangle light scattering (SEC-MALS) and circular dichroism (CD) measurements to uncover the varying oligomeric status and thermal stabilities. These reconstituted complexes were subjected to structural analysis using SAXS and electron microscopy, demonstrating the overall architecture of mammalian Nup88•Nup62 and Nup88•Nup62•Nup214 heterotrimers. Notably, we observed a significant similarity between the structures of Nup62•Nup88•Nup214 heterotrimer and earlier reported Nup62•Nup54•Nup58•Nup93 complex, which led us to establish that the Nup93 can bind to Nup62 shared compositionally different heterotrimers such as Nup88•Nup62, Nup88•Nup62•Nup214, Nup62•Nup54•Nup58 and Nup62•Nup54. Thus, our data here illuminate the hub role of Nup62 to form distinct heterotrimers and of Nup93, which behaves as a sensor in recognizing these Nup62 shared heterotrimers; thereby assembling the CR and CTC complexes of the mammalian NPC.

## RESULTS

### Divergence in the sequences of Nup88 complex across phyla

The phylogenetic analysis of Nup88, Nup62 and Nup214 from a total of 38 different organisms showed that Nups from human, rat and other vertebrates like *Xenopus, Danio,* etc clustered distantly from the branches of *Chaetomium* and *Saccharomyces* (Fig. S2 and Table S1). The metazoan clade marks close proximity and similarity amongst various organisms from different phyla. Accordingly, the secondary structure prediction of Nup88, Nup62 and Nup214 from different organisms ranging from fungi to human displays the differences and similarities in domain organisation of these Nups in different organisms (Fig. 1a-1c). It is observed that the α-helical region in the mammalian Nup214 is shortened when compared to unicellular eukaryotes (Fig. S3). In mammals, this helical region is sandwiched between the propeller domain and the FG region, whereas in fungi, it is present at the C-terminal end (Fig. 1c). As reported previously (Chopra et al 2019), we noticed that there are significant divergences in the primary sequence identity between the Nups of fungi and vertebrates (Table S2). Thus, we concluded that although some of the domains and folds remain conserved, there are distinct species specific features evident among these Nups’ homologs from unicellular eukaryotes to vertebrates.

### Establishing the interacting interface among Nup88, Nup214 and Nup62

Earlier, we developed a computational tool, CoRNeA (Co-evolution Random forest and Network Analysis), which can predict binary interacting interfaces solely based on the primary protein sequences (Chopra et al 2020). We employed this tool to predict precise interacting regions among Nup88, Nup214 and Nup62. A concatenated multiple sequence alignment file (only structured regions) of two interacting partners namely: Nup62•Nup88, Nup88•Nup214 and Nup214•Nup62 were used in the CoRNeA workflow as described previously (Chopra et al. 2020). The final interface outputs were sorted based on convolution scores as high scoring residue pairs indicating a very high probability of these residue pairs to form the interactive interface between the two proteins (Fig. S4). The summarised representation of such inter-domain interactions has been depicted in figure 2a. Overall we observed that the Nup62 coiled-coil region forms an extensive interaction network with the coiled-coil region of Nup88 and Nup214. Similarly, the β-propeller domains of Nup88 and Nup214 are capable of interacting with each other; however, fewer contacts were formed between Nup62 coiled-coil regions with the β-propellers of Nup214 and Nup88. The detailed pairwise interaction analysis is described as following:

**Figure 2:**
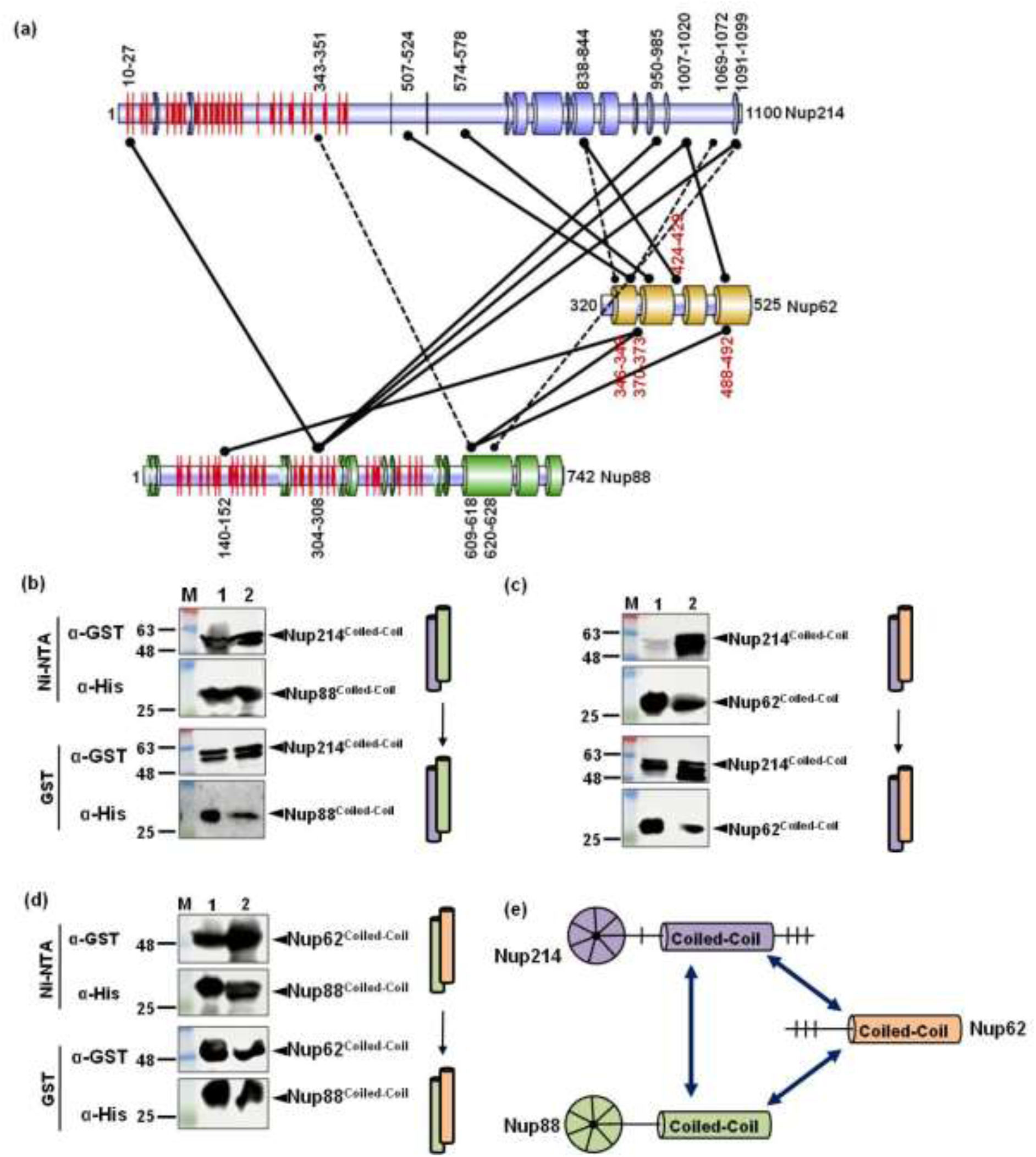
Interaction mapping among Nup88, Nup62 and Nup214 coiled-coil regions. (a) Schematic depiction of various inter-domain interactions between Nup88 (purple), Nup214 (green) and Nup62 (orange) by *in silico* tool CoRNeA. The helices are represented by cylinder and β-sheets by red colored rings. Interacting residue pairs are marked on the respective domains. High convolution score predictions are marked by solid lines while low score ones are represented by the dashed lines. (b-d) Western blot scan showing tandem affinity pull down (Ni-NTA followed by GST affinity) of coiled-coil domains (depicted as cylinders) of Nup88, Nup62 and Nup214. Lane M, 1 and 2 represent marker, input and eluent, respectively. Schematic representation of Nups88 (green) and Nup62 (orange) coiled-coil interactions as cylinders are shown. (b) GST tagged Nup214^693-976^ interaction with His_6_ tagged Nup88^517-742^. (c) GST tagged Nup214^693-976^ interaction with His_6_ tagged Nup62^322-525^. (d) His_6_ tagged Nup88^517-742^ association with GST tagged Nup62^322-525^. (e) Cartoon representation of summarising interaction results of CoRNeA prediction and pull down experiments.

#### a. Nup88 and Nup214 interface

For this pair, based on convolution scores (Fig. S4a), the strongest interacting regions were predicted for Nup88^β-propeller^ (304-308 amino acid residues) with Nup214^β-propeller^ (10-27) and Nup214^coiled-coil^ (950-1099 amino acid residues). Low scores were obtained for regions: Nup88^coiled-coil^ (611-628) interacting with Nup214^coiled-coil^ (1091-1100) and Nup214^β-propeller^ (201-351) (Fig. S4a). Overall, it indicated that both the β-propeller and coiled-coil domain of Nup214 and Nup88 can interact with each other.

#### b. Nup214 and Nup62 interface

CoRNeA analysis of this pair revealed the α-helical region of Nup62 (370-490) has strong interaction with Nup214^coiled-coil^ (685-1072). Very low convolution scores were obtained for Nup214^β-propeller^ with Nup62^coiled-coil^ indicating low possibility of direct interactions among them (Fig. S4b).

#### c. Nup88 and Nup62 interface

A very strong interaction between Nup62 and Nup88 was observed. A high convolution score suggest Nup62^coiled-coil^ (370-492) and Nup88^coiled-coil^ (609-626) have strong interaction between them. Interestingly, CoRNeA also predicted an interface for Nup62^coiled-coil^ (370-373) and Nup88^β-propeller^ (140-152) with high scores (Fig. S4c).

### Pull down approach delineates the inter-domain interaction network among Nup88, Nup62 and Nup214

To test the CoRNeA predicted domain interactions, we evaluated their *in vitro* binding capability using tandem affinity pull-down assay. In each case, we co-transformed the recombinant plasmids harbouring specific regions of two Nups fused with two different tags (GST and His_6_), respectively. Then we performed protein overexpression and subsequently tandem affinity purification (Ni-NTA pull-down in series with GST pull-down) to analyse their physical interaction with each other. Further, the interactions were confirmed by western blotting using anti-His_6_ and anti-GST antibodies, respectively.

#### a. Nup88coiled-coil interacts with Nup62coiled-coil and Nup214coiled-coil in a stable manner

GST tagged Nup214^coiled-coil^ (693-976) was co-transformed with either His_6_ tagged Nup62^coiled-coil^ (322-525) or Nup88^coiled-coil^ (517-742) for tandem affinity purification. We observed that the Nup214 coiled-coil region (693-976) has stable interaction with the coiled-coil regions of Nup88 and Nup62 (517-742 and 322-525, respectively) (Fig. 2b and 2c). In another parallel set of pull-down experiments, the GST tagged Nup62^coiled-coil^ region was co-expressed with the His_6_ tagged Nup88^coiled-coil^. They were also subjected to similar analysis, and physical interaction was seen between the coiled-coil regions of these two Nups (Fig. 2d). Thus, it became evident (Fig. 2e) that the coiled-coil domain of Nup88, Nup214 and Nup62 are capable of forming a stable complex as they are not washed away during the two steps of the purification. Furthermore, this analysis validated our *in silico* predictions (Fig 2a), where, we noticed a high convolution score for the coiled-coil domain of Nup62 and Nup88 and relatively lower convolution scores for Nup214^coiled-coil^ and Nup62^coiled-coil^ interface (Fig. S4).

#### b. Both β-propeller and α-helical domains of Nup88 and Nup214 interact with each other

The β-propeller domain of Nup88 (59-498) was co-expressed as His_6_-tagged fusion protein with either the helical or β-propeller domain of Nup214 *viz* Nup214 (1-407) and Nup214 (693-976) which were fused with the GST tag. They were purified by tandem affinity purification. We found that the β-propeller region of Nup88 was able to pull-down both the β-propeller and coiled-coil regions of Nup214, forming a probable α/β and α/α association, respectively. The interactions appear to be biochemically stable as clearly visible in western analyses of the co-eluting fractions (Fig. 3a and 3b). Similarly, in the case of Nup214 (59-498) co-expressing with Nup88 (517-742), we noticed that both proteins were interacting stably (Fig. 3c). Taken together these data and *in-silico* interface prediction data (Fig. 2a), we concluded that the β-propeller and coiled-coil domains of both Nup88 and Nup214 interact with each other making a lengthwise strong connection (Fig. 3d).

**Figure 3:**
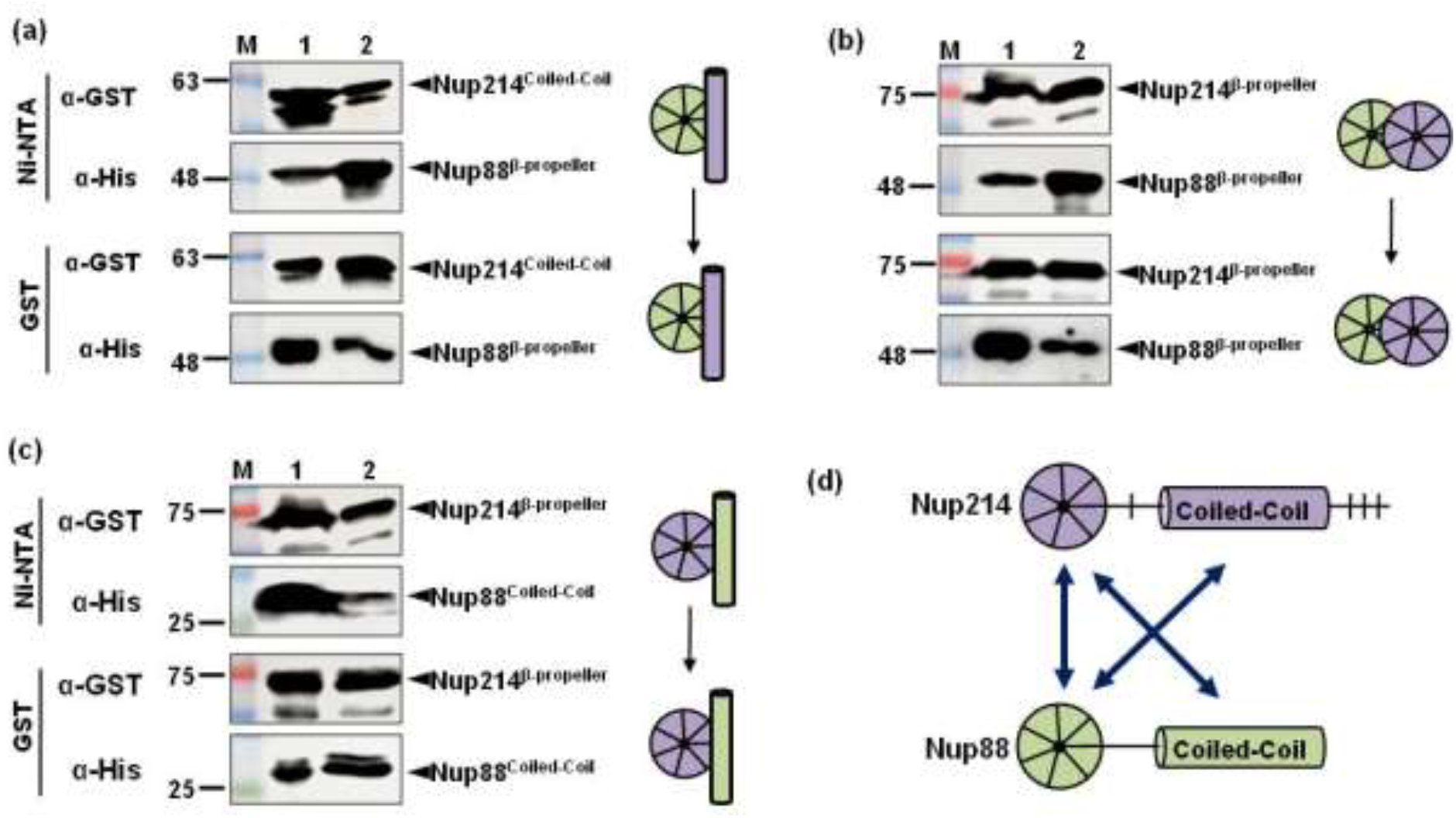
Interaction mapping of coiled-coil and β-propeller domains of Nup88 with Nup214. (a-c) Western blot scan showing tandem affinity pull down (Ni-NTA followed by GST affinity) of Nup88 (green) and Nup214 (purple). Lane M, 1 and 2 represent marker, input and eluent, respectively. Schematic representation of coiled-coils of Nup88 (green) and Nup62 (orange) and β-propeller domains are depicted as cylinders and circles, respectively. (a) GST tagged Nup214^693-976^ interaction with His_6_ tagged Nup88^59-498^. (b) GST tagged Nup214^1-407^ interaction with His_6_ tagged Nup88^59-498^. (c) GST tagged Nup214^1-407^ interaction with His_6_ tagged Nup88^517-742^. (d) Cartoon representation of summarising interaction results of CoRNeA prediction and pull down experiments.

#### c. The coiled-coil domain of Nup62 interacts transiently with β-propeller domain of Nup214 and Nup88

Nup88^β-propeller^ is the region between amino acid residues 59-498, while Nup214^β-propeller^ is between residues 1-407. When Nup62^coiled^ ^coil^ (322-525) co-elution with the β-propeller regions of Nup214 and Nup88 were analysed, we observed that the bands of His_6_-tagged Nup62^coiled-coil^ and Nup88^β-propeller^ were washed off in tandem GST purification (Fig. 4a and 4b) suggesting a relatively transient interaction between these domains (Fig. 4c). Similar weak or no interaction was also observed in our CoRNeA based prediction analysis (Fig 2a).

**Figure 4:**
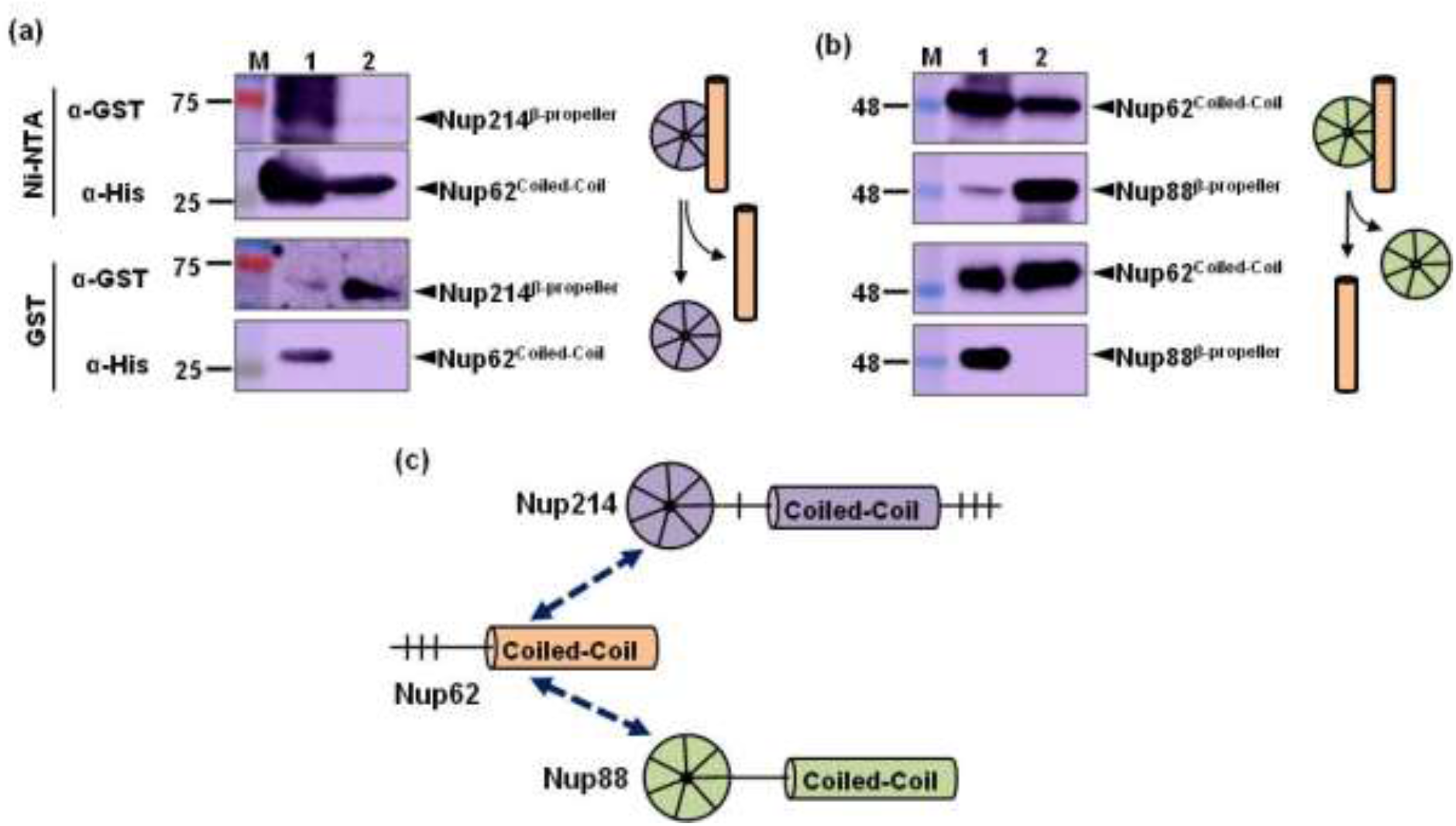
Mapping interaction of Nup62 coiled-coil domain with β-propeller domains of Nup88 and Nup214. (a-b) Western blot scan showing tandem affinity pull down (Ni-NTA followed by GST affinity) of Nup88 (green), Nup62 (orange) and Nup214 (purple). Lane M, 1 and 2 represent marker, input and eluent, respectively. Schematic representation of Nup88 and Nup214’s β-propeller domain are depicted as circles; and helical regions of Nup62 as a cylinder. (a) GST tagged Nup214^1-407^ showed transient interaction with His_6_ tagged Nup62^322-525^. (b) GST tagged Nup62^322-525^ has weak interaction with His_6_ tagged Nup88^59-498^. In both the cases, one of the partners is washed away during the second round of purification. (c) Cartoon representation of summarising interaction results of CoRNeA prediction and pull down experiments.

### α-helical domains are sufficient for Nup88•Nup62•Nup214 complex reconstitution

Based on our *in silico* and pull-down analyses, we established that the coiled-coil regions of Nup88, Nup62 and Nup214 are capable of interacting with each other. Additionally, to accomplish biochemical reconstitution of Nup88•Nup62•Nup214 assembly, we co-expressed and purified the helical regions of these Nups (Fig. 5a). The tandem affinity purified proteins (Fig 5b and 5c) were subjected to size exclusion chromatography (SEC) after fusion tag removal which indicated the formation of stable complex under given biochemical conditions (Fig 5d). However, we found ambiguity in validating the presence of all three Nups on SDS-PAGE as it showed the presence of two or/and three closely placed bands (Fig 5d). It seemed that all the three proteins (Nup88=26.6kDa; Nup62=23.7 kDa and Nup214=27.4 kDa) migrated very close to each other. Therefore, these proteins were further subjected to orbitrap mass spectrometry, where trypsin digested fragments of all the three Nups were detected as top hits (Fig. S5a and b and Table S3). Thus, the formation of trimeric complex (Nup88•Nup62•Nup214) by *in vitro* reconstitution was confirmed.

**Figure 5:**
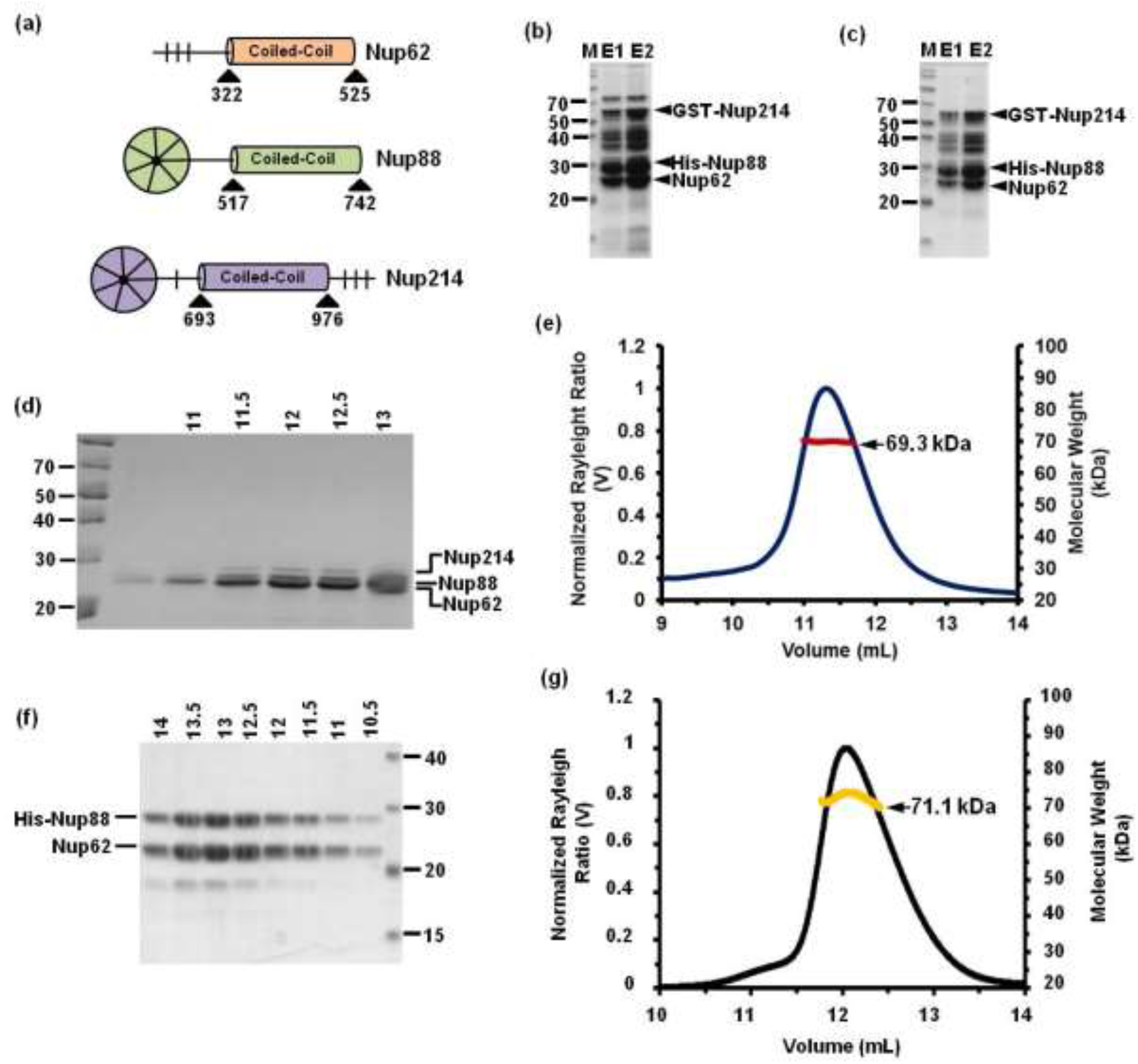
Purification of Nup88•Nup62 and Nup88•Nup62•Nup214 complexes. (a) Diagrammatic depiction of domain boundaries of Nup62, Nup88 and Nup214 used for cloning, co-expression and purification. (b) SDS-PAGE showing the purified heterotrimeric Nup88•Nup62•Nup214 complex via GST-affinity purification followed by (c) Ni-NTA affinity purification. (d) 12% SDS-PAGE scan showing the SEC peak eluted fractions (ml) of Nup88•Nup62•Nup214 complex. (e) SEC-MALS analysis of the Nup88•Nup62•Nup214 complex showing the Rayleigh scattering in black and molecular weight distribution across the peak in red color. (f) 12% SDS-PAGE scan showing the SEC eluted peak fractions (ml) of Nup88•Nup62 complex. (f) SEC-MALS analysis of the Nup88•Nup62 complex showing Rayleigh scattering in black while yellow color marks molecular weight distribution across the peak.

### Establishing the Oligomeric status of Nup88•Nup62•Nup214

We have earlier reported that the coiled-coil region of Nup62 (322-415) exists in a dynamic equilibrium between trimer and dimer (Dewangan, Sonawane et al. 2017). Further, the sequences of Nup88 and Nup214 when analyzed by MULTICOIL2 indicated that the propensity of these nucleoporins to form trimers is more than to form dimers. To understand the oligomeric properties of individual proteins we purified them separately as Nup62 (322-525) and Nup88 (517-742) (Fig. S6a and S6b). The purification of Nup214 (693-976) alone did not yield a soluble form of the protein (data not shown). We therefore concluded that Nup214 certainly needed an interacting partner to remain in a soluble form. Consistent with this MULTICOIL2 prediction, SEC-MALS analysis of Nup88 (517-742) (theoretical molar mass with tag is 26.6 kDa) indicated two peaks ranging from 75 kDa to 50 kDa suggesting a dynamic equilibrium between trimer and dimer, respectively (Fig S6c). Also, the peak for the trimer was much higher when compared to that of the dimer. Similarly, The purified Nup88^517-742^•Nup62^322-525^•Nup214^693-976^ complex, when analyzed using SEC-MALS, showed the average molar mass of 69.3 kDa (theoretical mass 77.5 kDa) (Fig. 5e), which suggests the complex is likely to be a hetero-trimer with one copy each of Nup88, Nup62 and Nup214. We used two different concentrations of the complex (0.4 mg/ml and 1.4 mg/ ml) to rule out the concentration dependent variations and found the averaged molar mass to be consistent (table S4). The stoichiometry of 1:1:1 was further confirmed by the glutaraldehyde cross linking exhibiting a protein band corresponding to 75 kDa (Fig. S7). Thus, the recombinantly expressed Nup88•Nup62•Nup214 complex is inferred to be a heterotrimer with 1:1:1 stoichiometry (Fig. 5e and table S4). In case of *X. laevis*, similar heterotrimer model was fitted into the EM density map which indicated the coiled-coils from three core proteins of the Nup88 complex (Huang, Zhang, Zhu, Zeng et al. 2020). Furthermore, our finding is similar to reported stoichiometry of corresponding homologous complexes in fungi. (Gaik, Dirk, Von Appen et al. 2015: Fischer, Teimer et al. 2015). This indicates that the propensity of these Nups to form heterotrimer is evolutionarily conserved.

### Nup88^517-742^ and Nup62^322-525^ forms a dynamic heterotrimer

Based on our protein-protein interaction analysis, we observed that the coiled-coil domains of Nup88 and Nup62 interact with each other strongly. The affinity purified Nup88•Nup62 complex, when analyzed using SEC methods, showed a homogeneously reconstituted complex (Fig 5f). Further, we obtained a uniform distribution of averaged mass (71.1 kDa) with SEC-MALS (Fig. 5g). Since the theoretical mass of the hetero-dimeric complex is 52.8 kDa, we suspect that the complex exists in 1:2 stoichiometries with an additional chain of either Nup62 or Nup88. To confirm this, we did a densitometric analysis of the SDS-PAGE bands, which clearly specify that the band intensity corresponding to the Nup62 is almost twice of the Nup88; indicating two copies of Nup62 in complex with single Nup88 (Fig. S8).

### Solution structure reveals an elongated shape of heterotrimer Nup88^517-742^•Nup6^2322-525^ complex

To gain further insight into the Nup88•Nup62 complex structure and dynamicity, we employed the small angle X-ray scattering (SAXS) technique, an established method for the structural analysis of dynamic behaviour of proteins in solution. We calculated the radius of gyration (R_g_) using the low Q region as well as the radius of cross section (R_c_) by assuming the globular and rod-like shape of the predominant scattering molecules in the solution. By employing the L= [12(R_g_^2^–R_c_^2^)]^1/2^ (Table 1), the length of persistence (L) of respective scattering entities was calculated. Further, the double log plot i.e. Log10 I (Q) versus Log10 Q (nm^-1^) confirmed no inter-particle interaction or aggregation (Fig. S9a). The Guinier analysis, Ln I (Q) versus Q^2^ plot is linear indicating good monodisperse quality of the heterodimeric complex (Fig. S9b). The slope of the Guinier plot was used to calculate the R_g_ value of the complex as 7.11 nm (Table 1). The D_max_ was calculated from the pairwise distance distribution plot P(R) using indirect Fourier transformation and the maximum distance observed is 36.39 nm. The elongated and highly dynamic shape of the P(R) functions confirms the extended state of the complex in solution. The Kratky plot i.e. I (Q)*Q^2^ versus Q (nm^-1^) shows the highly flexible structure of the complex corroborated by Porod exponent (quantitative metric for assessing compactness) value of 2.2 (Fig. S9c and S9d) suggesting a less compact and highly dynamic structure.

**Table 1:**
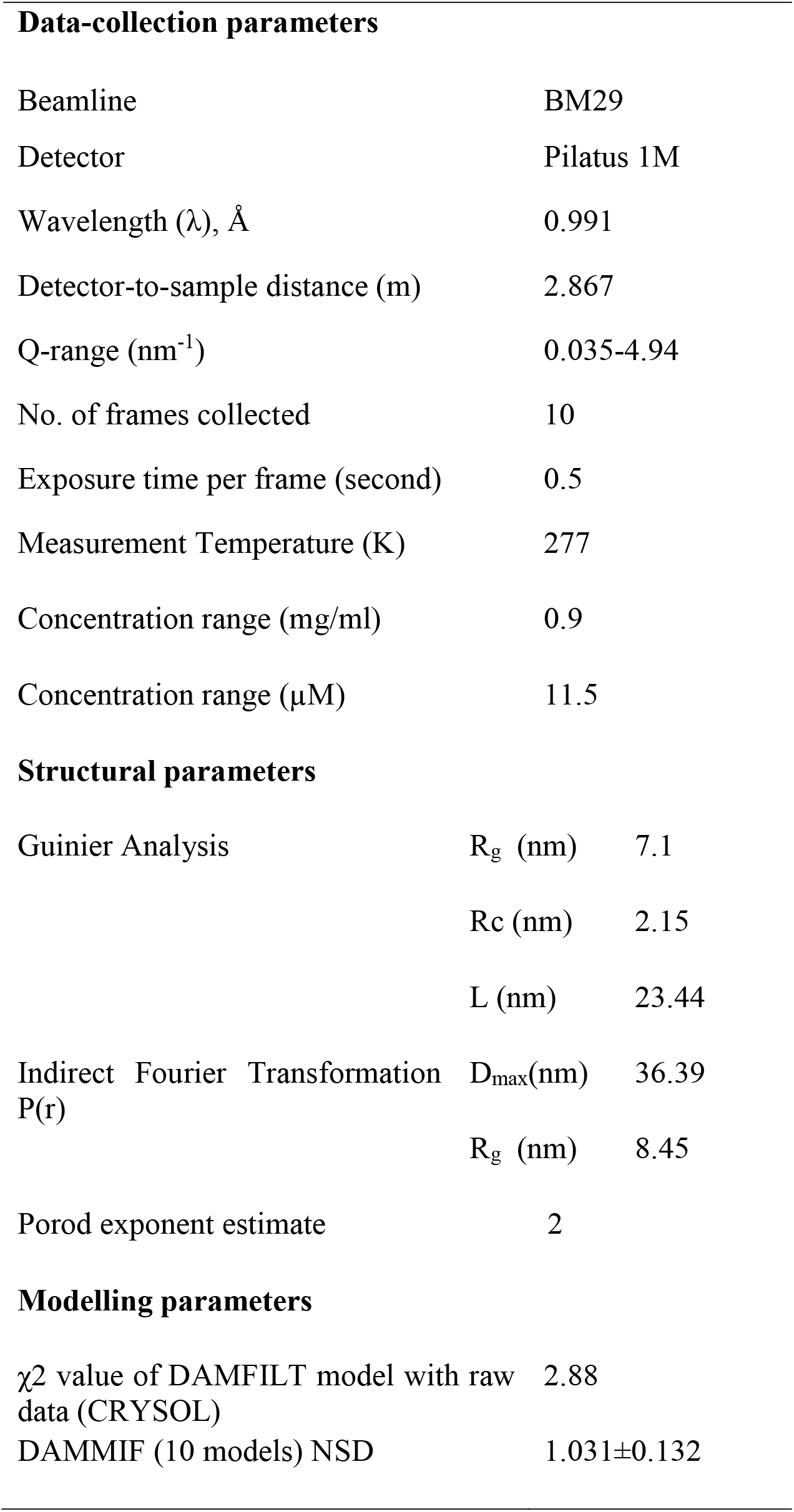
SAXS data collection, structure solution, and model parameters for Nup88•Nup62 complex.

| Data-collection parameters |  |  |
| --- | --- | --- |
| Beamline | BM29 |  |
| Detector | Pilatus 1M |  |
| Wavelength ($\lambda$), Å | 0.991 | |
| Detector-to-sample distance (m) | 2.867 |  |
| Q-range (nm<sup>-1</sup>) | 0.035-4.94 |  |
| No. of frames collected | 10 |  |
| Exposure time per frame (second) | 0.5 |  |
| Measurement Temperature (K) | 277 |  |
| Concentration range (mg/ml) | 0.9 |  |
| Concentration range (µM) | 11.5 |  |
| Structural parameters |  |  |
| Guinier Analysis | R<sub>g</sub> (nm) | 7.1 |
|  | R<sub>c</sub> (nm) | 2.15 |
|  | L (nm) | 23.44 |
| Indirect Fourier Transformation P(r) | D<sub>max</sub>(nm) | 36.39 |
|  | R<sub>g</sub> (nm) | 8.45 |
| Porod exponent estimate | 2 |  |
| Modelling parameters |  |  |
| χ<sup>2</sup> value of DAMFILT model with raw data (CRY SOL) | 2.88 |  |
| DAMMIF (10 models) NSD | 1.031±0.132 |  |

Based on the SAXS data, we performed dummy atomic modelling (DAMMIF) to build the 3D model of the complex. The final generated model of the complex was compared with the raw data I (Q) profile by using the CRYSOL program. The Chi-square value of 2.88 suggests that the DAMFILT model fits well with the raw data. There are some extra regions observed at the extremities of the SAXS model which could be due to intrinsic flexibility of the protein complex in solution (Fig. 6). High degree of flexibility between the coiled-coil domains of fungal homolog, Nup82 with a kinked structure was also reported earlier (Fernandez-martinez, Kim, Shi, Upla, Gagnon, Pellarin et al. 2016). Additionally, we earlier showed that the coiled-coil domain of Nup62 forms a triple helix bundle (Dewangan, Sonawane et al. 2017) and our SEC-MALS data in this study, further suggests that the Nup88•Nup62 complex exists in the 1:2 stoichiometry. Hence, we represented Nup88•Nup62 as a heterotrimer in 1:2 stoichiometry as shown in figure 6. It appears that the complex is highly elongated. Overall, our SAXS data clearly indicates that Nup88•Nup62 complex possesses a flexible and dynamic behaviour in solution and adopts an elongated structure.

**Figure 6:**
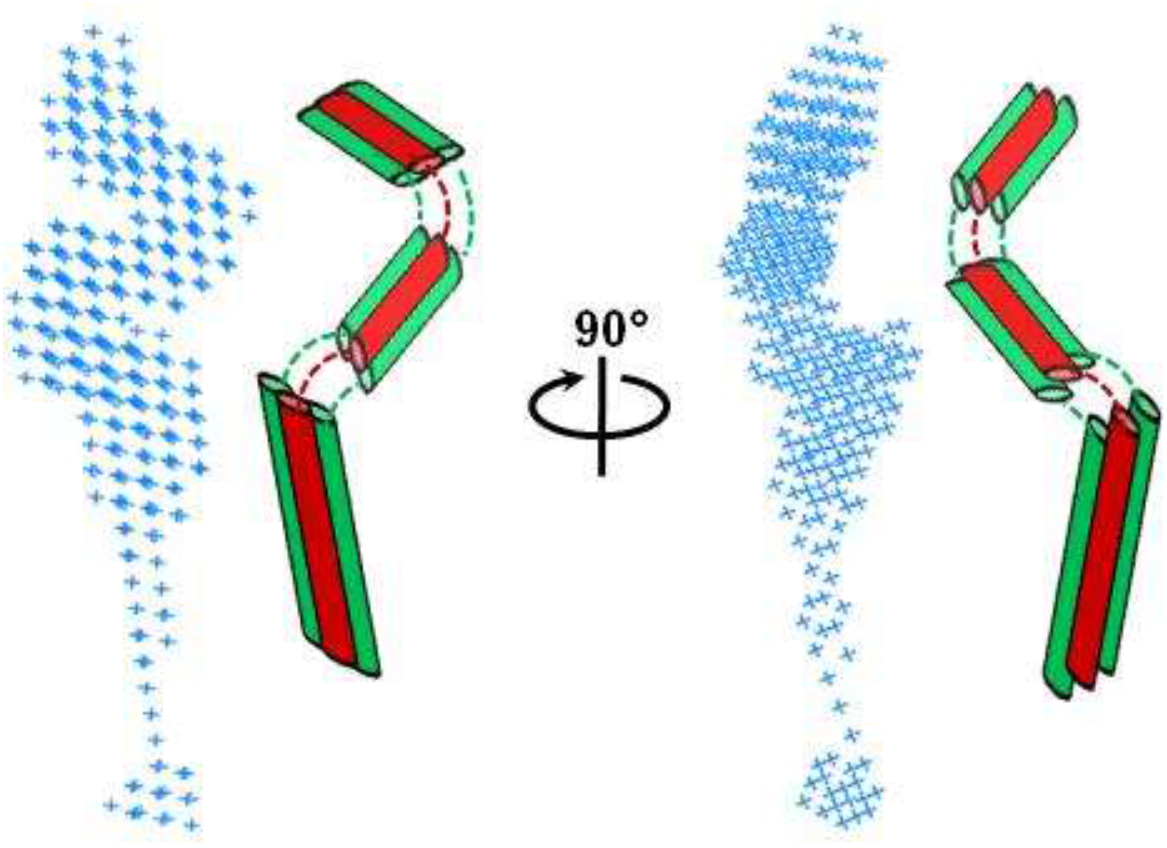
Small Angle X-ray Scattering (SAXS) analysis of Nup88^517-742^•Nup62^322-525^ complex. Molecular envelope of Nup88^517-742^•Nup62^322-525^ complex obtained by DAMMIF analysis (shown in blue color). The cartoon model of the protein complex (two copies of Nup62 in green color and one copy of Nup88 in red) is shown in two different orientations.

### Nup88•Nup62 and Nup88•Nup62•Nu214 complexes show variable conformational thermostability

The α-helical, coiled-coil is one of the principal subunit oligomerization motifs in proteins. Despite its simplicity, it is a highly versatile folding motif due to which coiled-coil-containing proteins exhibit a broad range of functions (Greenfield and Hitchcock 1993; Burkhard et al. 2001). We probed the secondary structure and conformational stability of proteins by circular dichroism (CD) spectroscopy. Wavelength scans showed the α-helical coiled-coil signature with minima at 208 nm and 222 nm for Nup62 and Nup88 (Fig. S6d), Nup62•Nup88 complex and Nup62•Nup88•Nup214 complex (Fig. 7a). The *θ*_222_/ *θ*_208_ ratio depicts the helix propensity of the proteins. Also, the combination of helix propensity and hydrophobic core packing determines the stability of coiled-coil structures (Patricia et al. 2019). So, we calculated the *θ*_222_/ *θ*_208_ ratios and % helicity in each case as displayed in the table S5a. When the *θ*_222_/ *θ*_208_ ratio exceeds 1, it indicated a well-defined coiled-coil structure. It is clear from the table S5a that the complexes behave very well as coiled-coil structures in solution.

**Figure 7:**
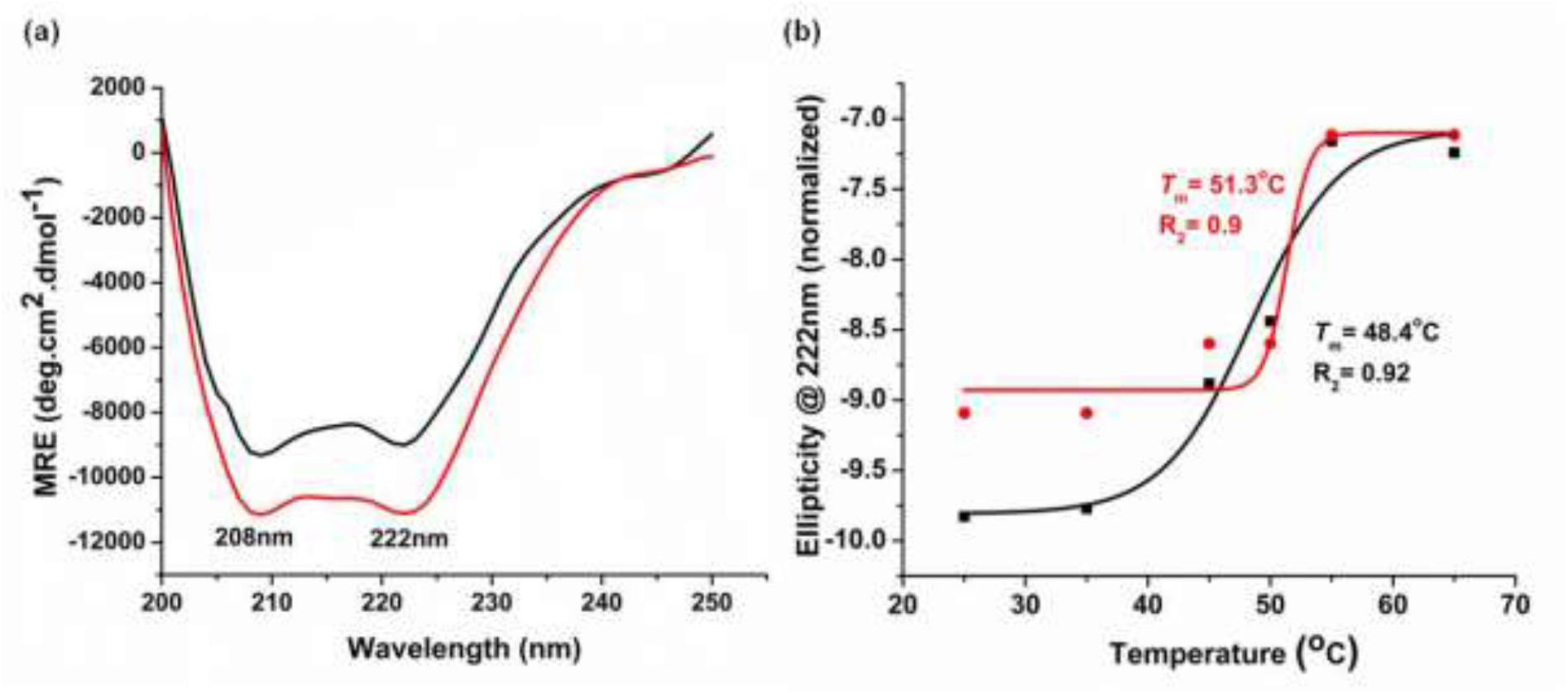
CD spectroscopy analysis of the Nup88•Nup62 and Nup88•Nup62•Nup214 complexes. (a) Far-UV CD spectra of purified Nup88•Nup62 and Nup88•Nup62•Nup214 are shown in black and red colour, respectively. (b) Thermal denaturation profile showing *T*_m_ for the complexes via sigmoidal fit of the ellipticity shift at 222 nm. The red colour denotes curve for the Nup88•Nup62•Nup214 complex while the curve for Nup88•Nup62 is shown in black color.

To compare the thermostability of the heterodimeric and heterotrimeric complexes, we investigated their thermal denaturation with CD spectroscopy. Examination of the CD signal at 222 nm over a range of temperatures (25°C–95°C) showed that the Nup88•Nup62 complex exhibits *T*_m_ of 48°C and the Nup88•Nup62•Nup214 complexes have a *T*_m_ of 51°C (Fig. 7b), a feature that is consistent with the thermostability of other naturally occurring coiled-coil domains (*T*_m_ ≥ 50°C) (Fujiwara et al. 2012; Tsuruda 2011). It is apparent from the CD denaturation curves that though the unfolding begins, the Nup88•Nup62•Nup214 complex retains its helical structure even at higher temperatures (40-45°C). On the other hand, the Nup88•Nup62 complex loses its helical conformation faster and is adopting random coil structure as the temperature increases (Table S5b). These results suggest that Nup214^693-976^ plays an important role in providing stability to the complex. Moreover, the thermal denaturation was irreversible in all the cases (data not shown).

### Structure of the Nup88•Nup62•Nup214 complex revealed by negative stain electron microscopy (EM)

As we could not pursue SAXS studies of Nup88•Nup62•Nup214 complex due to the low solubility, we analysed the structure of this heterotrimer using negative stain electron microscopy. The purified complex was adsorbed on EM grids and stained with 2% Uranyl acetate. Approximately, 4737 particles were selected and utilized for the single particle analysis using CryoSPARC (Fig. 8a). The 2D class averages obtained showed the distinct “elongated structures with a curve at one end” (Fig. 8b). Further, these 2D classes were used for the ab-initio 3D reconstruction and refinement (Fig. 8c). Based on the FSC cut-off at 0.143, the global resolution of the structure is 19.27Å (Fig. S10b). In comparison with SAXS data of Nup88•Nup62 complex (Fig. 6) we observed that, by adding the Nup214 into the heterotrimer, the complex is a more compacted and stable structure (Fig. 8c) which is also reflected by the thermal stability analysis (Fig 7b). The overall architecture of the Nup88•Nup62•Nup214 complex showed an elongated ‘4’ shape with the bulky head. This shape shows similarity with the tube-like density reported for the coiled-coil helix bundle of Nup88 complex in *Xenopus* (Huang, Zhang, Zhu, Zeng et al. 2020) where they modelled full length Nup88 and the coiled-coil domains of Nup62 and Nup214 and fitted into the tomogram as cryo-ET reconstruction. We obtained the homology models of the Nup88, Nup62 and Nup214 from the AlphaFold database (Varadi et al. 2022) and built a complex model using the *Ct*Nsp1 complex (PDB ID; 5CWS; Stuwe, Bley, Thierbach, Petrovic et al. 2016) as the template. This modelled complex was then docked into the EM density map (Figure 8d), which shows that the model fitted very well in the density map and adopts a ‘4’ shape structure inside the density, which is almost identical to reported *Ct*Nsp1•Nup49•Nup54•Nic96 complex (Stuwe, Bley, Thierbach, Petrovic et al. 2016); where about ~41 residues (139-180) of Nic96 (a homolog of Nup93) is located in the bulky head. Moreover, our previous study (Sonawane, Dewangan et al. 2020) have shown that mammalian Nup93 (1-150) region is capable to form a quaternary complex with CTC (Nup62•Nup54•Nup58) and form a ‘4’ shaped structure where the CTC complex is aligned parallel to Nup93. Interestingly a feature of ‘bulky head’ was also observed.

**Figure 8:**
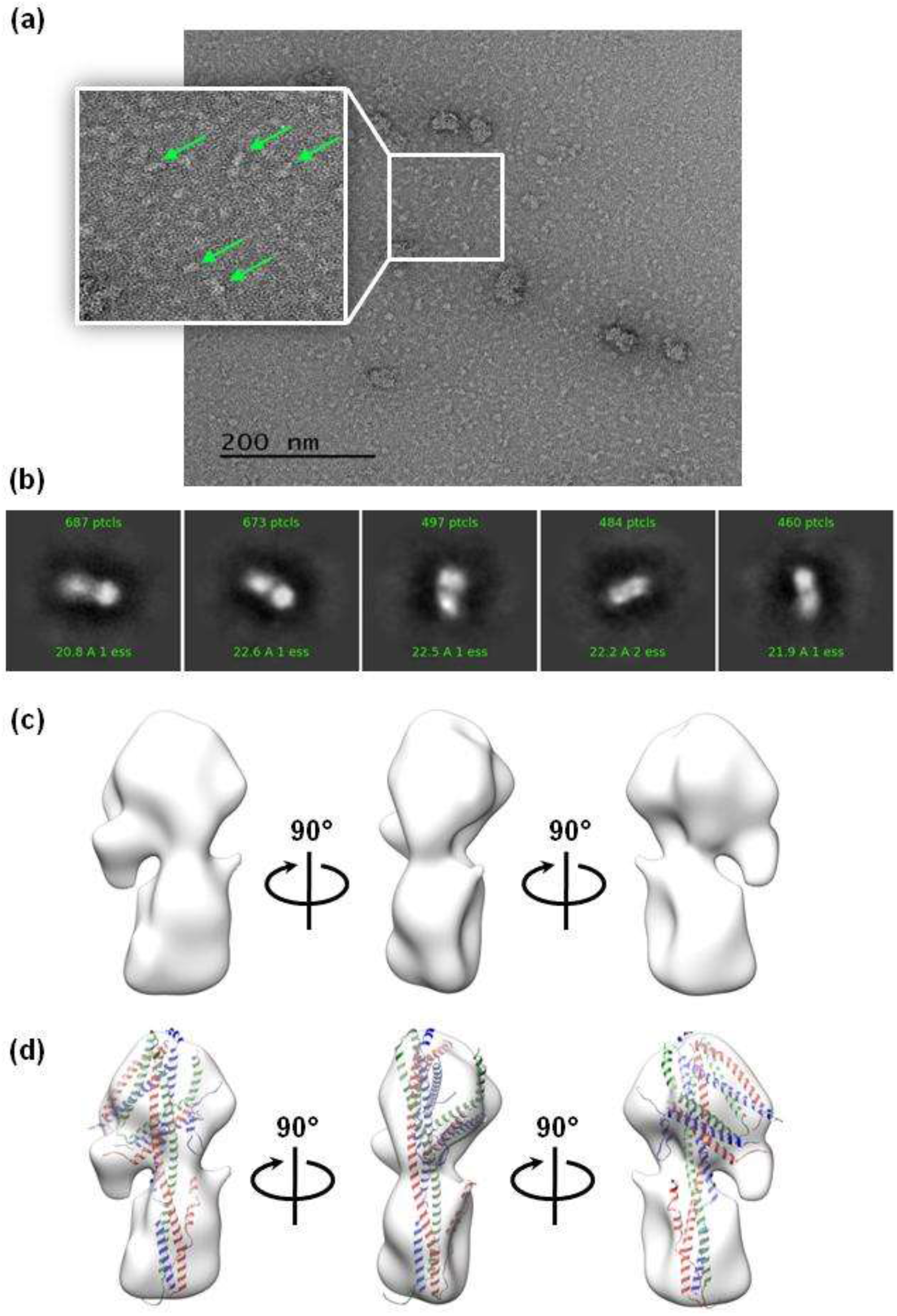
Negative staining EM analysis of Nup88^517-742^•Nup62^322-525^•Nup214^693-976^ complex. (a) A representative micrograph of purified Nup88•Nup62•Nup214 complex stained with 2% uranyl acetate. Scale bar: 200 nm. Representative particles used for the analysis are shown in inset with the arrow. (b) Representative 2D class averages. (c) 3D density map obtained at 19.27 Å. (d) The density map fitted with modelled structures of the coiled-coil complex.

### Nup88•Nup62 and Nup88•Nup62•Nup214 complex interacts with Nup93^1-150^

The superimposition of the EM density of CTC•Nup93 (1-819) complex with the density map obtained for the Nup88 complex (Fig. S10 c-e) revealed that, the head region of both the densities matches perfectly indicating the similar arrangement of triple helix bundle, formed by both Nup88•Nup62•Nup214 and Nup62•Nup58•Nup54. So, based on these observation, we hypothesised that similar to CTC complex, the CR complex of Nup88•Nup62•Nup214 may also bind to the Nup93 and we evaluated if the Nup93^1-150^ (known to form stable complex with CTC; Sonawane, Dewangan et al. 2020) can interact with either or both Nup88•Nup62 and Nup88•Nup62•Nup214 heterotrimers. We performed tandem affinity purification of both these complexes with the Nup93 (1-150) domain. To our surprise, we found a stable interaction of Nup93^1-150^ not only with Nup88•Nup62•Nup214 (Fig. 9 a-b) complex but also with Nup88•Nup62 heterotrimer (Fig. 9 c-d). This clearly establish that highly dynamic Nup88•Nup62 complex attains a similar heterotrimeric conformation to provide interface for Nup93^1-150^ region similar to Nup88•Nup62•Nup214 heterotrimer.

**Figure 9:**
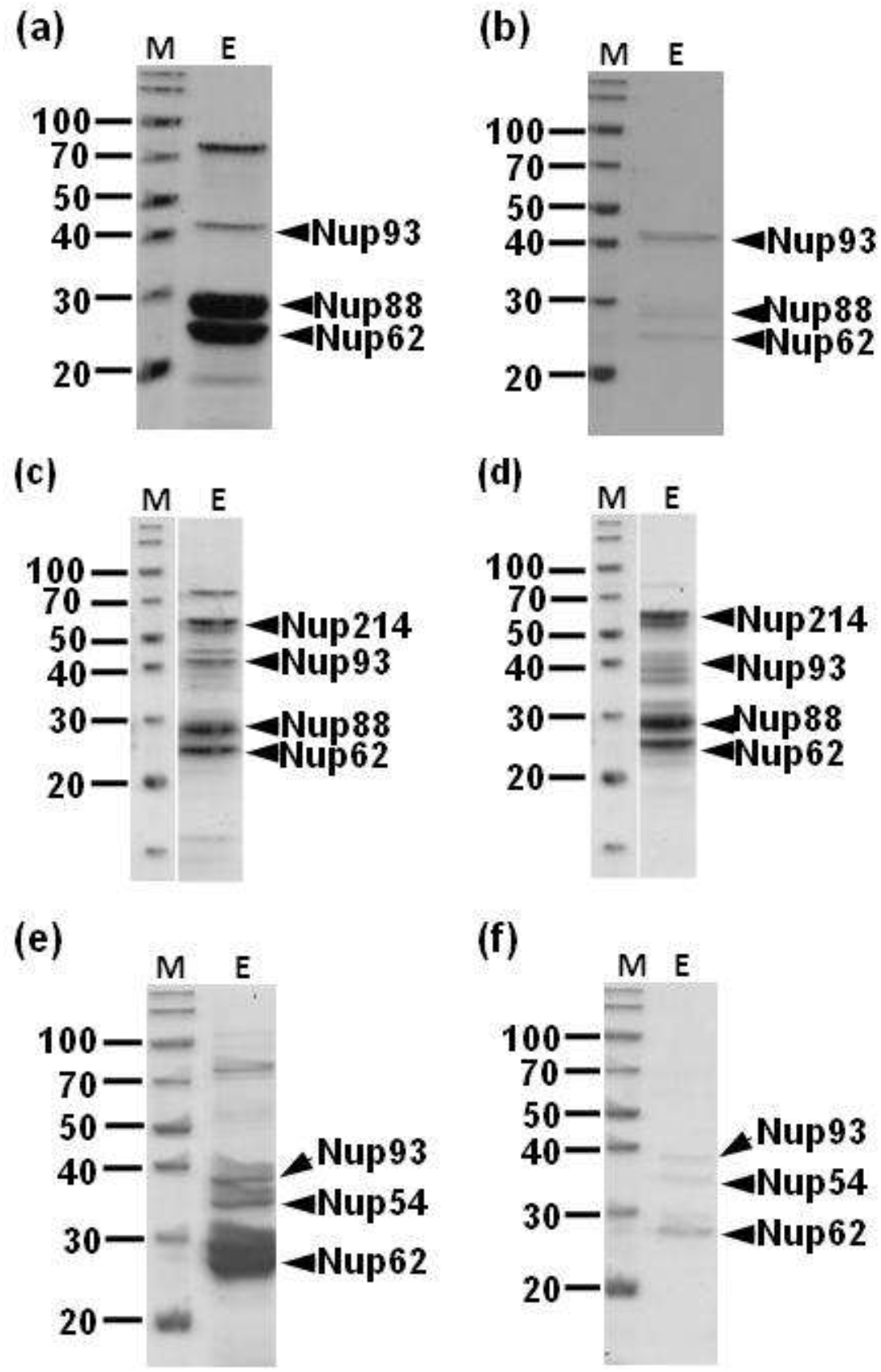
Interaction analysis of N-terminal domain (1-150) of Nup93 with Nup88•Nup62, Nup88•Nup62•Nup214 and Nup62•Nup54 complexes. SDS-PAGE images showing the reconstituted complexes with Nup93^1-150^ and their interaction via tandem affinity purification (Ni-NTA affinity followed by GST affinity purification). (a and b) Nup88•Nup62 complex. (c and d) Nup88•Nup62•Nup214 complex. (e and f) Nup62•Nup54 complex. a, c and e: Ni-NTA affinity elutions; b,d and f: GST-affinity elutions. Lane 1 and 2 in each image correspond to protein marker and elution, respectively.

### Mammalian Nup62•Nup54 heterotrimer also can bind with Nup93

Previously it was reported that Nup62 along with Nup54 can form a stable heterotrimer (Solmaz, Radha et al. 2011) and its minimal coiled-coil domain structure was revealed using X-ray crystallography showing 2 helices of Nup62 interacting with one helix of Nup54. Interestingly, in this case also, it is observed that mammalian Nup62•Nup54•Nup58 heterotrimer is highly dynamic and Nup58 can be easily dissociated to form Nup62•Nup54 complex (Sharma et al. 2015). We found such observation remarkably similar to our reported data in case of Nup88•Nup62 complexes. Therefore, we made an assumption that similar to Nup88•Nup62 heterotrimer, Nup62•Nup54 heterotrimer may bind to the Nup93^1-150^ region. To prove this, we co-transformed a plasmid containing human Nup54 containing all structured regions (181-507) and Nup93^1-150^ encoding genes with rat Nup62 (322-525) into *E. coli* cells and performed a tandem affinity pull down experiment as described earlier. The SDS-PAGE analysis clearly indicated that mammalian Nup62•Nup54 heterotrimer is also capable of interacting with Nup93^1-150^ in a stable manner as it did not dissociate during the tandem affinity purification steps (Fig. 9 e-f). The presence of the three proteins was also confirmed in the western analysis using anti-His_6_, anti-GST and anti-Nup54 antibodies (Fig. S11). This led us to conclude that Nup93^1-150^ region is capable to bind with compositionally different yet structurally similar heterotrimers such as Nup88•Nup62, Nup88•Nup62•Nup214, Nup62•Nup54 (described in this study) and Nup62•Nup54•Nup58 (described in Sonawane, Dewangan et al. 2020). Such phenomenon of multipartner protein interface clusters which can adaptively bind to different proteins due to their conserved binding pocket is also known in other cases in the literature (Keskin and Nussinov 2007).

## DISCUSSION

The CR of the mammalian NPC is an important platform for mRNA export and is linked to several diseases. To gain insights into the inter-domain interactions and structural aspects, we focused on the mammalian Nup88•Nup62•Nup214 complex positioned in the CR along with the Y-shape complex. In this study, we show for the first time, extensive interaction network and biochemical reconstitution of the mammalian Nup88•Nup62 and Nup88•Nup62•Nup214 complex. We observed that most of the interacting regions are conserved between mammalian and corresponding complexes in fungal species (Gaik, Dirk, Von Appen et al. 2015; Teimer et al. 2017) indicating that the conserved core assembly of the CR complex is driven by the coiled-coil domains of the Nup62, Nup88 and Nup214. The Nup62 is also a key component to form the CTC complex as reported previously (Bailer et al. 2000; Solmaz, Chauhan et al. 2011) in similar stoichiometry and molecular arrangement, it is therefore reasonable to propose that the Nup62 coiled-coil domain, which is highly dynamic in nature (Dewangan, Sonawane et al 2017), play an important role in providing the plasticity of both CR and CTC complex.

The biochemical reconstitution followed by characterization using CD spectroscopy, SAXS, EM, and SEC-MALS analysis of Nup88•Nup62•Nup214 and Nup88•Nup62 heterotrimers established in solution, the dynamic behaviour of Nup62•Nup88 complex to form a highly elongated complex. Our data also show that incorporation of the Nup214 into Nup88•Nup62 formed a slightly more stable heterotrimer that results in a compact conformation (Fig. 8d). The low-resolution 3D structure of the Nup88•Nup62•Nup214 complex by negatively stained electron microscopy showed elongated density with a bulky and curved head. It displayed an unusual asymmetric ‘4’ shaped structure where helical regions of all three Nups can be modelled. As our interaction analysis demonstrated direct physical interaction of β-propeller domains of Nup88 and Nup214 (Fig. 3b), they both are expected to occupy the bottom of the Nup88•Nup62•Nup214 complex (Fig. 10a).

**Figure 10:**
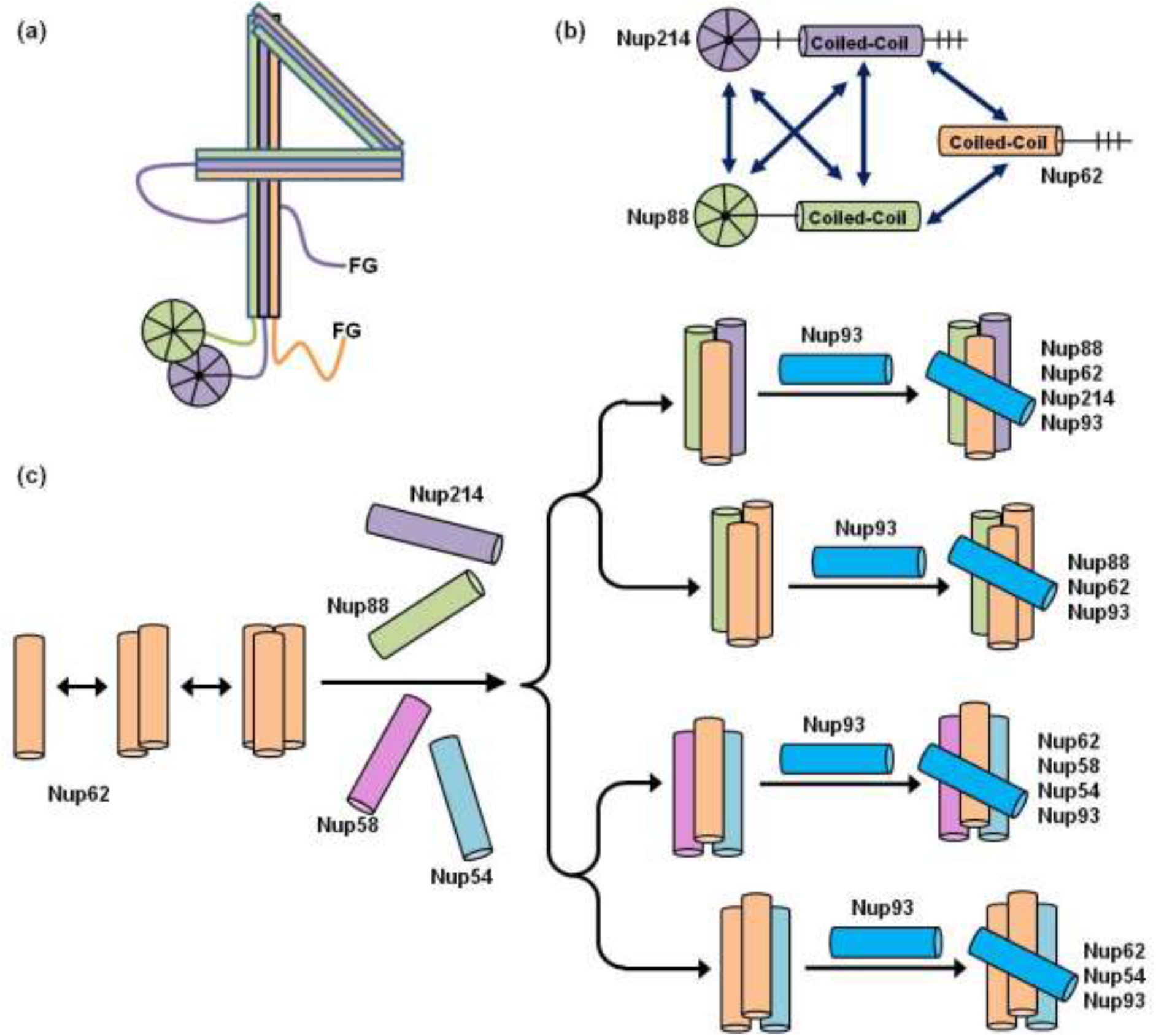
Role of Nup62 and Nup93 as hub proteins in the mammalian NPC. (a) Cartoon representation of ‘4’ shape structure adopted by the coiled-coil domains of the Nup88•Nup62•Nup214 complex and showing β-propeller domain interaction of Nup88 and Nup214. (b) Summary of the inter-domain interaction network among the three Nups. (c) Cartoon representation of coiled-coil domain of Nup62 (orange), Nup88 (green), Nup214 (purple), Nup54 (cyan) and Nup58 (pink). The plasticity exhibited by Nup62 coiled-coil domain allows it to interact with other Nups and form various heterotrimers such as Nup88•Nup62, Nup88•Nup62•Nup214, Nup62•Nup54 and Nup62•Nup54•Nup58. Such heterotrimers are further recognized by the N-terminal (1-150) of Nup93 (blue) to form distinct quaternary complexes.

We have previously reported the low resolution 3D structure of mammalian Nup62•Nup54•Nup58•Nup93 quaternary complex (Sonawane, Dewangan et al. 2020), and we observed remarkable similarity with Nup88•Nup62•Nup214 heterotrimer 3D map as reported here. Hence, we conclude that both complexes adopt a similar architectural arrangement of coiled-coils. Notably, it has been reported that in *C. thermophilum,* the Nsp1•Nup49•Nup57 complex (homolog of Nup62•Nup58•Nup54) is structurally related to the Nup82•Nup159•Nsp1 complex (situated in CR), as both complexes assume a similar triple helix coiled-coil ‘4’ shaped architecture (Chug et al. 2015; Teimer et al. 2017; Fernandez-Martinez, Kim, Shi, Upla, Gagnon, Pellarin et al. 2017). Such structural similarity is further demonstrated in our data where we showed that Nup93 (1-150) can bind to Nup88•Nup62•Nup214 heterotrimers (Fig. 9) in similar fashion as reported previously for Nup62•Nup54•Nup58 complex (Sonawane, Dewangan et al. 2020). Based on all these evidences, we propose that Nup62 is a hub protein of the NPC to form distinct biochemically stable heterotrimers (Nup62•Nup88•Nup214, Nup88•Nup62, Nup62•Nup54 and Nup62•Nup54•Nup58 complexes) and this ability is exhibited due to dynamic behaviour of its coiled-coil domain (Fig.10c).

These distinct heterotrimers which share Nup62 are shown to interact with the N-terminal domain of Nup93 (1-150) to form the quaternary complex (Fig. 9 a-f). Nup93 is central to the IR complex, which can interact with Nup188, Nup35, Nup205 and Nup155 (Fischer, Teimer et al. 2015). Additionally, it is shown to anchor the CTC complex previously indicating the role of Nup93 as a linker Nup (Fischer, Teimer et al. 2015). It is also reported previously that the N-terminal region of Nup93 (1-150) has two short helices (1-82 and 96-150) that can preferentially interact with other partners such as the region 1-82 interact with the CTC complex in 1:1:1:1 stoichiometry (Sonawane, Dewangan et al. 2020). Here, we show that the N-terminal region of Nup93 (1-150) can bind to a diverse set of heterotrimers formed by the Nup62. Moreover, in case of both fungal and mammalian CTC, so far only Nup62•Nup54•Nup58 complex is shown to interact with Nup93 (Sonawane, Dewangan et al. 2020; Stuwe, Bley, Thierbach, Petrovic et al. 2016). Although it is reported earlier that Nup62•Nup54 also forms a heterotrimer but its relevance with Nup93 interaction was not clear. Based on our interpretation of Nup93 mode of recognition of a heterotrimer, we could demonstrate that even Nup62•Nup54 heterotrimer is capable of interacting with Nup93. It is therefore plausible that Nup62 shares various heterotrimers that pose a specific binding pocket that can be recognized by Nup93. Such shared organisational behaviour of Nups is very intriguing in terms of NPC assembly and its versatile transport function.

In case of multipartner protein, a hub protein is defined as a protein with the conserved domain/motif that can interact with multiple partners (Ekman et al. 2006) and show highly clustered functional modules. In the case of Nup93 and Nup62, it is shown that both of these are essential Nups for cell viability. Role of Nup93 in NPC assembly and role of Nup62 in mRNA export and mitosis is extensively studied (Fernandez-martinez, Kim, Shi, Upla, Gagnon, Pellarin et al. 2017; Galy et al. 2003; Hashizume, Moyori et al. 2013; Okazaki, Yamazoe et al. 2020; Sachdev et al. 2010). Moreover, their functions are evolutionarily conserved. Our study here suggests that Nup93 and Nup62 meet these criteria to be hub proteins and thus enable us to undertake further studies to refine their role in the transport activity in a Nup specific manner.

Interestingly, though the overall structure of the Nup88 complex and its homolog in yeast (Nup82 complex) is similar; but, their interactions with other Nups could be different. For instance, the mammalian Nup88 complex interacts with the N-terminal of Nup93 as described here. However, in fungi, corresponding homolog Nup82 complex does not bind to the N-terminal of Nic96 (homolog of Nup93); rather it interacts with Nup145C (Teimer et al. 2017; unpublished data, Bley, Liu Nie, Mobbs, Petrovic, Gres et al. 2021). This highlights the fact that although both CTC and Nup88/82 complexes are conserved between unicellular and vertebrate NPC, their anchorage to the NPC framework might be mediated by two different partners, namely Nup145C and Nup93, respectively. It will be interesting to understand further how these differences would lead to species specific arrangement of IR and CR.

The human CR Nups and their associated mRNA export factors are recognized as important players in multiple disease conditions (Lin and Hoelz 2019; Yarbrough et al. 2013; Hurt and Alwin 2010; Nofrini et al. 2016; Ciomperlik al. 2016) and are suggested to govern the remodelling of mRNPs at the cytoplasmic face of the NPC (Bailer et al. 2000; Hutten and Kehlenbach 2006; Kalverda, Pickersgill et al. 2010; Napetschnig 2007; Snay-hodge et al. 1998). Therefore, the ultimate mechanistic insights of these Nups will be feasible by high-resolution structural analysis, which could not be pursued for mammalian Nup88 complexes till now, mainly due to the lack of biochemical reconstitution. Our interaction and reconstitution data here suggested a hub role for Nup62 in assembling both Nup88 complex (constituent of CR) and CTC complexes and the role of Nup93 in their anchorage to the NPC framework, thus laying the ground for atomic resolution structural studies in future.

## MATERIALS AND METHODS

### Sequence analysis of Nups

A total of 38 sequences were obtained from Uniprot (Table S1) for each Nup and were aligned by MUltiple Sequence Comparison by Log Expectation (MUSCLE). The sequences were assembled to a neighbour-joining tree using MEGA 6 (Tamura et al. 2013) and the bootstrap was calculated with 1000 replications. The final trees were edited and viewed using iTOL tool (Letunic and Bork 2019). The sequence alignment (structure guided) of rat, human, *Xenopus*, *Danio*, *Chaetomium*, and *Saccharomyces* was done using PROMALS3D (Pei et al. 2008) and edited in Jalview (Waterhouse et al. 2009) to remove the low complexity FG repeat regions and Jpred (Drozdetskiy et al. 2015) to predict the secondary structures considering *Rattus norvegicus* Nup as a template. The domain organisations based on the secondary structure predictions for the Nups were performed using the PSIPRED server (Jones 1999) and the sequences were visualised using the IBS illustrator (Liu et al. 2015).

### CoRNeA analysis

Co-evolution Random forest and Network analysis was performed using primary amino acid sequences from both the proteins in a complex as the input information as described in Chopra et al. 2020. Only the structured regions of Nup88 (1-742), Nup62 (320-525) and Nup214 (1-1100) were considered for the predictions.

### Generation of *E. coli* based over expression plasmids

The total RNA was isolated from rat spleen using Trizol reagent (Invitrogen) and cDNA was prepared by Superscript-II synthesis kit (Invitrogen) following the manufacturer’s instructions. The gene specific primers (Table S6) for Nup88, Nup62 and Nup214 were used to amplify various deletion constructs using Phusion^®^ polymerase (NEB). The α-helical of Nup62 and α-helical as well as β-propeller regions of Nup88 and Nup214 were cloned into the respective vectors (refer Table S6). In all the cases, affinity tags were removed by thrombin digestion. All positive clones were confirmed by gene sequencing.

### In vitro pull-down assay

pET28a and pGEX4T1 vectors having respective genes (Table S6) were co-transformed into *E. coli* BL21 (DE3) RIL cells. The bacterial culture was grown at 37°C until OD_600_ reached 0.6, induced with 0.4 mM of IPTG (MP Biomedicals) and incubated for 14-16 h at 18°C. The bacterial cells were harvested and lysed in lysis buffer (20 mM Tris-HCl pH 8, 250 mM NaCl, 1 mM PMSF, 5 mM β-Me, 10 mM imidazole, 10% Glycerol and 1% Triton X-100). The clear lysate was incubated with Ni-NTA agarose beads (Qiagen) for 2 h at 4°C. The unbound fractions were discarded and beads were washed with 40 column volume (CV) of wash buffer (20 mM Tris-HCl pH 8, 250 mM NaCl, 5 mM β-Me, 35 mM imidazole, 2.5% Glycerol and 0.1% Triton X-100). Bound fractions were eluted with elution buffer (20 mM Tris-HCl pH 8, 250 mM NaCl, 5 mM β-Me, 300 mM imidazole, 2.5% Glycerol and 0.05% Triton X-100). Ni-NTA eluted fractions were incubated with GSH-Sepharose (Pierce) beads for 5 h at 4°C. The unbound fractions were discarded and beads were washed with 40 CV of wash buffer (with 0.05% Triton X-100). Bound fractions were eluted with elution buffer (with 10 mM reduced glutathione). The eluted fractions from each pull down (Ni-NTA and GST-affinity) were probed with anti-GST (1:3000) antibody (Sigma) and anti-His (1:3000) antibody (Sigma). HRP conjugated mouse IgG was used at 1:5000 dilution (Sigma) for developing the signal and all the images were captured using Amersham^TM^ Imager 600 and 800 (GE Healthcare).

### Protein purification

To purify the Nup88•Nup62•Nup214 complex, pRSFDuet1 vector expressing Nup88 (517-742) and Nup62 (322-525) was co-transformed with pGEX4T1 expressing Nup214 (693-976) into the BL21 (RIL) cells. The protein was over expressed as described above and lysed using the lysis buffer mentioned previously followed by the centrifugation. The clear lysate was incubated with Ni-NTA (Qiagen) beads for 2-3 hrs at 4°C and washed with a 40X column volume wash buffer followed by elution. For the 2nd affinity step, the Ni-NTA eluted fractions were pooled, dialyzed and incubated with GSH-Sepharose (Pierce) beads. The bound complexes were eluted with a wash buffer having 10 mM of reduced glutathione (Sigma). Eluted fractions were the concentrated via 10 kDa concentrator (Amicon Ultra, Merck) and both His_6_ and GST tags were cleaved with 5-10 unit thrombin (Merck) per milligram of protein for 24 h at 4°C and subjected to the SEC using the superdex 200 10/30 GL column, pre-equilibrated with SEC-buffer (20 mM Tris-HCl pH8, 150 mM NaCl, 0.5 mM EDTA and 1mM DTT). The entire purification was analysed using SDS-PAGE. In order to purify Nup88•Nup62 binary complex the pRSFDuet-1 expressing Nup88 (517-742) and Nup62 (322-525) was similarly over expressed and purified as mentioned above. For the purification of Nup88 (517-742) and Nup62 (322-525), pET28a constructs were used and proteins were purified using Ni-NTA affinity chromatography. The buffer composition for Nup62 was similar except the detergent and glycerol, while for Nup88, 0.05% detergent was used. In order to purify Nup88•Nup62•Nup214•Nup93 complex, the bacterial cells having Nup88•Nup62•Nup214 and Nup93 (1-150) were expressed separately and co-lysed; purified by tandem affinity pull-down. For Nup88•Nup62•Nup93 complex, Nup88•Nup62 and Nup93 (1-150) were co-expressed and purified by tandem affinity pull-down. The buffer compositions were the same as mentioned above. The constructs for Nup54•Nup93 and Nup62 were co-transformed, over-expressed and purified using Ni-NTA followed by GST pull-down as described earlier (Sonawane, Dewangan et al. 2020).

### Mass spectrometry-based confirmation of Nup88•Nup62•Nup214 complex

The SEC eluted fraction and SDS-PAGE bands corresponding to the complex were subjected to trypsin digestion and investigated using Orbitrap analyzer and the obtained data were analyzed using Proteome Discoverer 2.2 (Thermofisher) software.

### Size exclusion chromatography coupled with multi angle light scattering (SEC-MALS)

The purified individual Nups and complexes at various concentrations (Table S4) were analysed with SEC using either Superdex 200 10/30 GL or Superose 6 increase 10/30 GL column coupled with multi-angle light scattering (MALS) instrument (Wyatt technology). The chromatography system was connected in series with a light scattering detector (Wyatt Dawn HELIOS II) and refractive index detector (Wyatt Optilab t-rEX). 2mg/mL of BSA (Sigma) was used as a standard to calibrate the system and 100µL of each sample was injected. The column was equilibrated with the SEC buffer. Data analysis was carried out using the program ASTRA (Wyatt Technologies), yielding the molar mass and mass distribution (polydispersity) of the samples.

### Circular Dichroism (CD) analysis and thermal denaturation

CD spectra of SEC purified proteins (0.15-0.2 mg/mL) were recorded using a Jasco J-815 (Jasco, Tokyo, Japan) spectropolarimeter at 25 °C. The proteins in 10 mM Tris-Cl (pH 8.0), 50 mM NaCl and 1 mM DTT used for Far-UV CD spectra were recording in a rectangular quartz cuvette of 1 mm path length in the range of 200–250 nm at a scan speed of 100 nm/min with slit width of 1 nm. Each spectrum was recorded as an average of 3 accumulations and was corrected for buffer contributions before analysing. The observed values were converted to mean residue ellipticity (MRE) using the equation: MRE = Mθ_λ_/10dcr, where *M* is the molecular weight of the protein, *θ_λ_* is CD in millidegree, *d* is the path length in cm, *c* is the protein concentration in mg/mL, and *r* is the average number of amino acid residues in the protein. The relative content of secondary structure elements was calculated using CDPro software and CONTINLL program which gave the least NRMSD (normalized root mean square deviation) value. The percent helicity (*α*-helix content) was calculated from the MRE value at 222 nm, using the equation: % α-helix = (MRE_222nm_ /MRE_max_) × 100, where MRE_max_ = −23400 (Wang et al. 2006). For thermal denaturation studies, the protein sample was incubated at varying temperatures from 25 to 85°C with an interval of 5°C for 5 min each. The melting temperatures of the proteins/complexes were calculated by scattering the ellipticity at 208 nm and 222 nm, followed by sigmoidal fit analysis.

### Small angle X-ray scattering (SAXS) analysis

Purified Nup88•Nup62 complex (0.9 mg/mL) was used to collect scattering data at European Synchrotron Radiation Facility (ESRF), Grenoble, France on beamline BM29 (λ=0.991 Å) in a standard setup (Pernot et al. 2013). Total 10 frames were collected (0.5s/frame) at 277K on PILATUS 1M detector and sample to detector distance was 2.87 mm. Sample buffer (SEC buffer) was used as control for buffer scattering datasets subtraction automatically via EDNA (Bourenkov et al. 2009). Scattering datasets collected were analysed using ATSAS 2.8 (Franke et al. 2017). Initial molecular envelope was built using DAMMIF. CRYSOL was used to obtain the theoretical I (Q) profile of the model and compared it with I (Q) profile of raw data.

### Negative staining and Electron microscopy

The 4 µL of the purified protein complex (0.05 mg/mL) was adsorbed onto a glow discharged carbon coated 300 mesh copper grid for 2 min and the excess sample was blotted away using Whatman^TM^ filter paper. Then, 4 µL of 2% uranyl acetate adjusted to pH 7.4 was applied on the grid twice for 2 min each. The grids were again blotted to remove excess stain and air dried overnight. The grids were loaded on JEM 220FS (Jeol) accelerated at 200 kV with a field emission gun. 54 micrographs were collected and subjected to single particle analysis using CryoSPARC (Punjani et al. 2017) using MotionCorr2 and CTF estimation was performed using CTFFIND4 implemented in RELION 3.1.0 (Jasenko et al. 2018). After several rounds of 2D classification followed by manual inspection, a total 4737 particles were selected to perform *ab-initio* reconstruction. The homogenous refinement was performed using a 3D map to obtain the final resolution.

## ACKNOWLEDGMENTS

We thank Dr. Srikanth Rapole, NCCS Mass Spectrometry Facility for help with MS analysis. The authors thank members of the Laboratory of Structural Biology, NCCS, for their critical inputs. PKJ and JS thank the Department of Biotechnology, India and University Grants Commission, India, respectively for the senior research fellowship. This work was supported by the Centre of Excellence in Biomolecular Structure and Function on Host-Pathogen Interactions grant (RC/PR/15450) to RC and NCCS intra-mural funding to RC. Data collection at the ESRF-SAXS BM29 beamline was facilitated by the ESRF Access Program of RCB, supported by the Department of Biotechnology, Government of India.

## Author contribution statement

RC designed the study. PKM, ES, JS, SB and MYA performed experiments and contributed in writing. PKM, ES and RC wrote the manuscript.

