## supplemental figures for "Insights into the role of Nup62 and Nup93 in assembling cytoplasmic ring and central transport channel of the nuclear pore complex"

### SUPPLEMENTARY INFORMATION

#### Figure S1

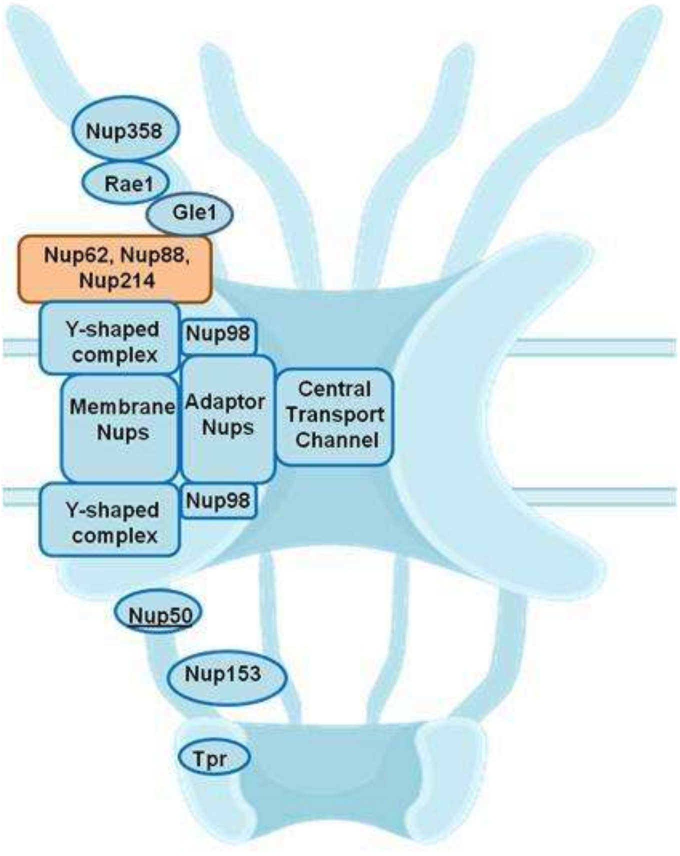


**Fig. S1: Schematic architecture of the NPC:** The cylindrical symmetric core of NPC is decorated with cytoplasmic ring and filaments and a nuclear basket. Natively unfolded FG repeats present in one-third of Nups make up the transport barrier in the central channel. The location of Nup88 complex is highlighted in the CR.

#### Figure S2

##### a) Nup88

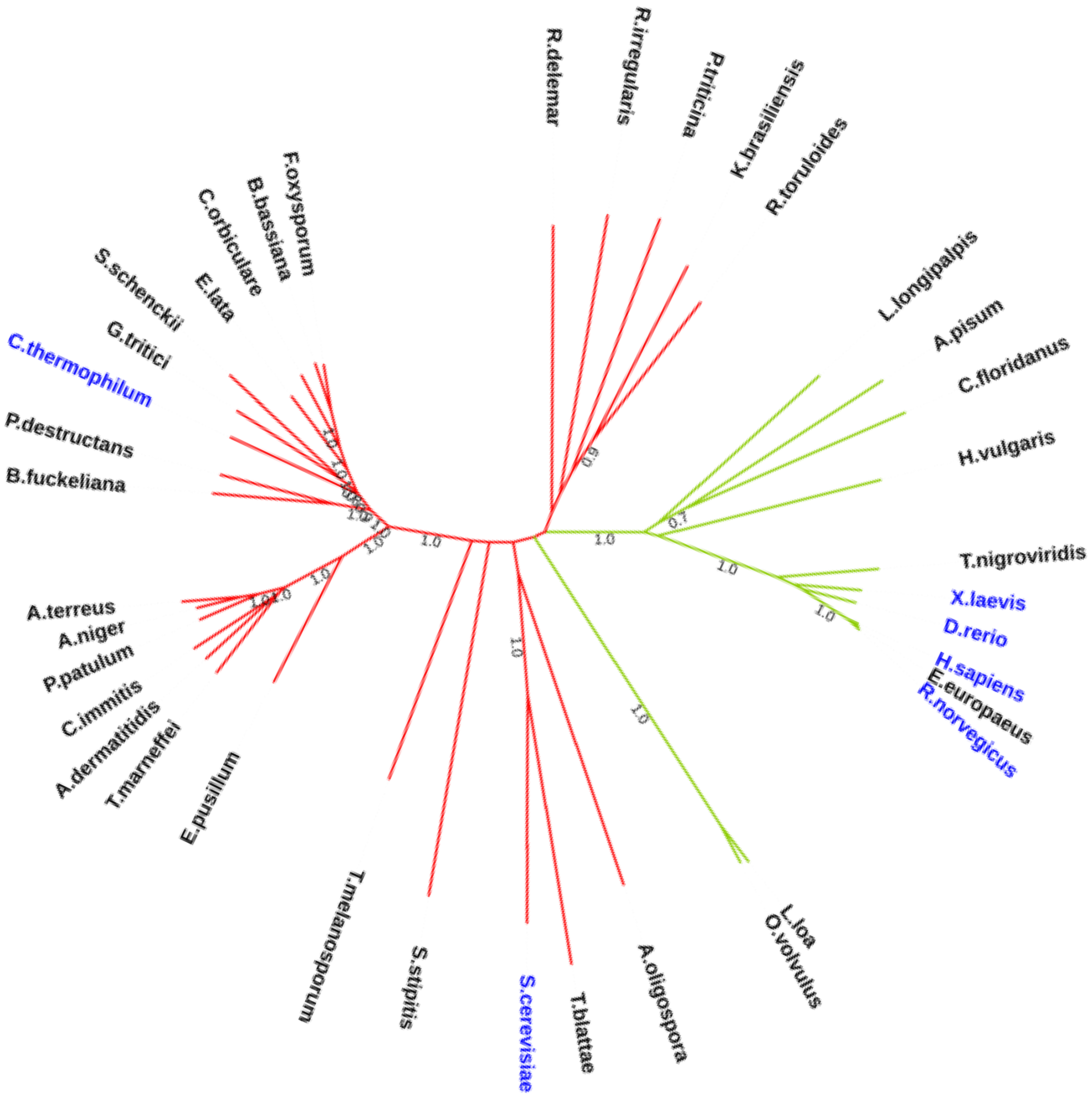

**b) Nup62**

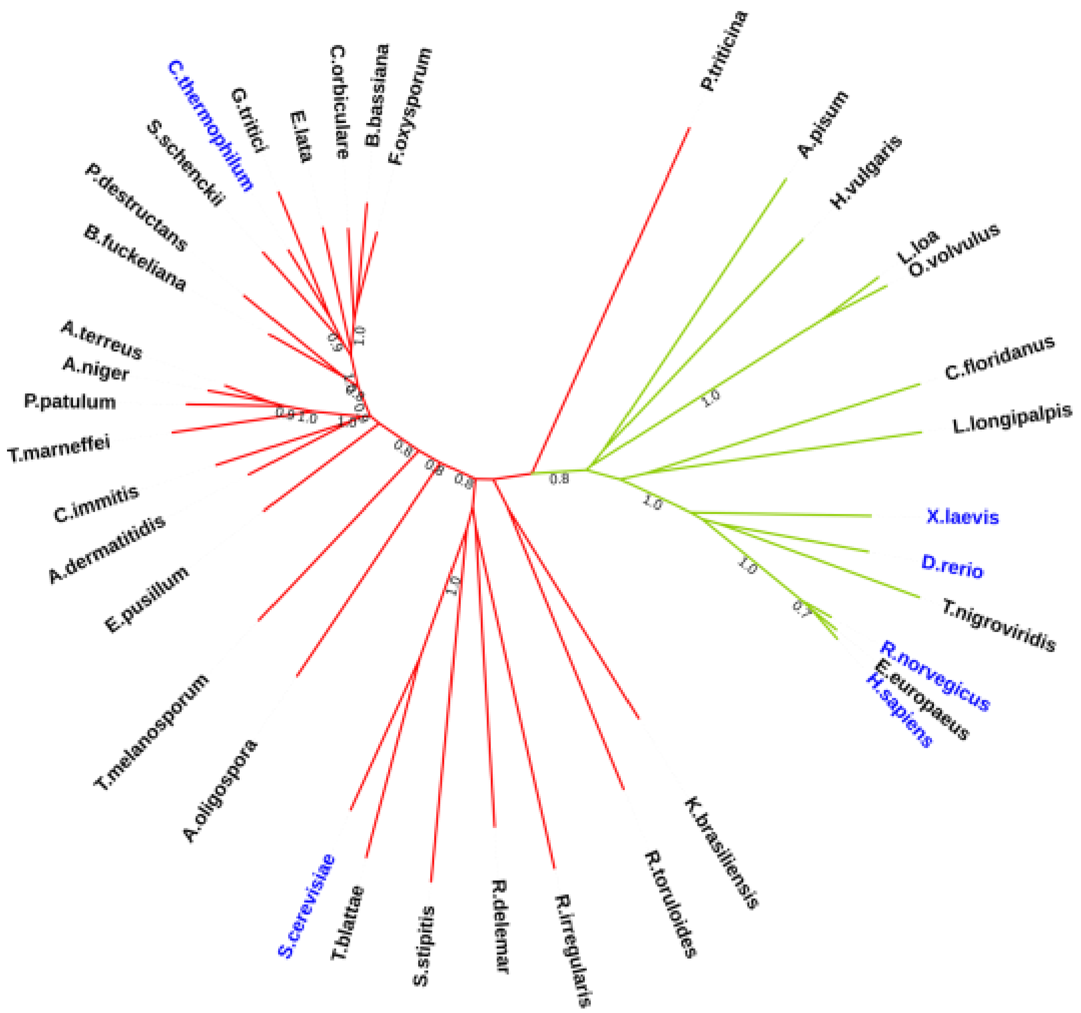
P.triticina
A.pisum
H.vulgaris
L.loa
O.volvulus
C.floridanus
L.longipalpis
X.laevis
D.rerio
T.nigroviridis
R.norvegicus
E.europaeus
H.sapiens
K.brasiliensis
R.toruloides
R.irregularis
R.delemar
S.stipitis
T.blattae
S.cerevisiae
A.oligospora
T.melanosporum
E.pusillum
A.dermatitidis
C.immitis
T.marneffei
P.patulum
A.niger
A.terreus
B.fuckeliana
P.destructans
S.schenckii
C.thermophilum
G.tritici
E.lata
C.orbiculare
B.bassiana
F.oxysporum

c) <u>Nup214</u>

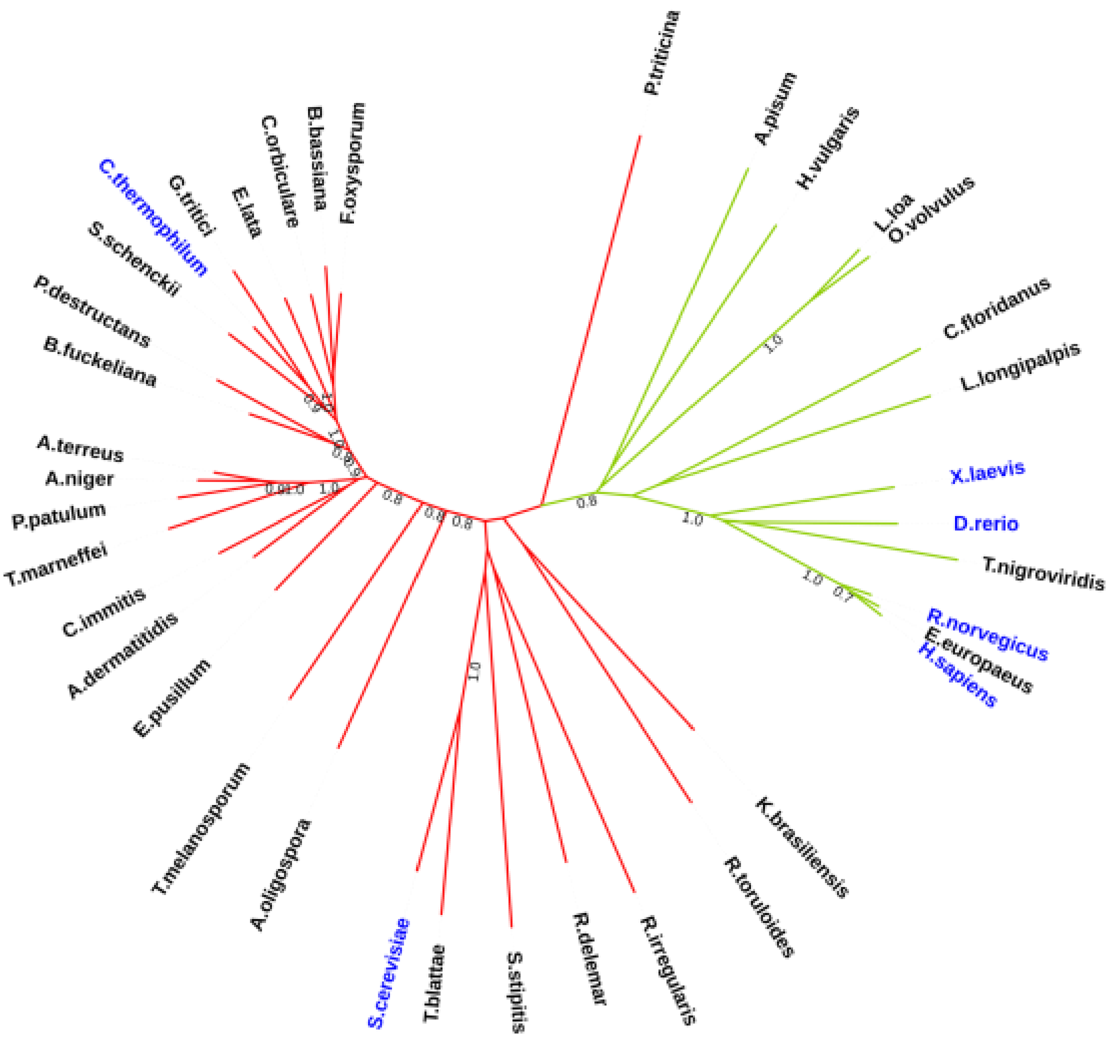


**Fig. S2: Phylogenetic analysis for homologs of (a) Nup62 and (b) Nup214** (related to Fig.1)**:** Unrooted phylogenetic trees were constructed using 38 homologs from different species across phyla. Only those species were chosen for which homologs of all three Nups (Nup88, Nup62 and Nup214) are reported in UniProt. The numbers on each branch represent bootstrap value and branches in green color highlights the metazoan clade. The Nup sequences for species shown in blue color have been compared in figure 1 and S3.

**Table S1: Details of species and sequences used for construction of phylogenetic trees for Nup88, Nup62 and Nup214**

| Sr. No. | Species | Classification | Uniprot Id | | |
|---|---|---|---|---|---|
| | | | Nup88/ Nup82 | Nup62/ Nsp1 | Nup214/ Nup159 |
| 1 | *Acyrthosiphon pisum* | Arthropoda | J9JUX4 | J9JW91 | J9JP64 |
| 2 | *Ajellomyces dermatitidis* | Ascomycetes | F2TFB1 | F2TMQ1 | F2T4C8 |
| 3 | *Arthrobotrys oligospora* | Ascomycetes | G1X4K3 | G1XR62 | G1XGH1 |
| 4 | *Aspergillus niger* | Ascomycetes | G1X4K3 | A0A3F3RGM3 | A0A100IJC4 |
| 5 | *Aspergillus terreus* | Ascomycetes | A0A5M3ZC47 | A0A5M3ZFR7 | A0A5M3Z4Q9 |
| 6 | *Beauveria bassiana* | Ascomycetes | A0A2N6P1K2 | A0A2S7Y588 | A0A2N6NM17 |
| 7 | *Botryotinia fuckeliana* | Ascomycetes | G2YSN1 | G2YTP6 | G2YZM4 |
| 8 | *Camponotus floridanus* | Arthropoda | E2A5P2* | E2AU26 | E2AUB7 |
| 9 | *Chaetomium thermophilum* | Ascomycetes | G0S4F3 | G0SBQ3 | G0SBS8 |
| 10 | *Coccidioides immitis* | Ascomycetes | J3KHV4 | J3KDP8 | J3KID1 |
| 11 | *Colletotrichum orbiculare* | Ascomycetes | N4VBK1 | A0A484FI75 | N4W5E4 |

| | | | | | |
|---|---|---|---|---|---|
| 12 | *Danio rerio* | Chordata | A2CEI4 | E9QIQ3 | E7F5M0 |
| 13 | *Endocarpon pusillum* | Ascomycetes | U1G970 | U1HIL0 | U1HTA8 |
| 14 | *Erinaceus europaeus* | Chordata | A0A1S2ZLS2 | A0A1S3AJS5 | A0A1S3WKH3 |
| 15 | *Eutypa lata* | Ascomycetes | M7SGG9 | M7SYK5 | M7TDE7 |
| 16 | *Fusarium oxysporum* | Ascomycetes | A0A559KT89 | A0A559LWN6 | A0A559L6V9 |
| 17 | *Gaeumannomyces tritici* | Ascomycetes | J3P7J8 | J3NK05 | J3NRV8 |
| 18 | *Homo sapiens* | Chordata | Q99567 | P37198 | P35658 |
| 19 | *Hydra vulgaris* | Cnidaria | T2M6Q5* | T2MGF0 | T2MG31* |
| 20 | *Kalmanozyma brasiliensis* | Basidiomycetes | V5F0U6 | V5GS41 | V5GK17 |
| 21 | *Loa loa* | Nematoda | A0A1I7W3V2 | A0A1I7VW80 | A0A1I7W224 |
| 22 | *Lutzomyia longipalpis* | Arthropoda | A0A1B0CE61 | A0A1B0CB33 | A0A1B0CV73 |
| 23 | *Onchocerca volvulus* | Nematoda | A0A044R9K4 | A0A044SAJ1 | A0A044SPY2 |
| 24 | *Penicillium patulum* | Ascomycetes | A0A135M0B4 | A0A135LDW2 | A0A135LGZ9 |
| 25 | *Scheffersomyces stipitis* | Ascomycetes | A3LRG4 | A3LN95 | A3GEU8 |
| 26 | *Pseudogymnoascus destructans* | Ascomycetes | L8FS73 | L8FZJ9 | L8FUV8 |

| | | | | | |
|---|---|---|---|---|---|
| 27 | *Puccinia triticina* | Basidiomycetes | A0A180GMP3 | A0A180GV19 | A0A0C4F0W7 |
| 28 | *Rattus norvegicus* | Chordata | O08658 | P17955 | D4ACK1* |
| 29 | *Rhizophagus irregularis* | Mucoromycota | U9T9T5 | U9TSV8 | U9UPA7 |
| 30 | *Rhizopus delemar* | Mucoromycota | I1BU21 | I1C636 | I1CIE2 |
| 31 | *Rhodosporidium toruloides* | Basidiomycetes | M7X0A4 | M7WSS4 | M7X5G0 |
| 32 | *Sporothrix schenckii* | Ascomycetes | A0A0F2MCG4 | A0A0F2MKY4 | A0A0F2M365 |
| 33 | *Talaromyces marneffei* | Ascomycetes | A0A093V5G4 | A0A093Y3M4 | A0A093VL94 |
| 34 | *Tetrapisispora blattae* | Ascomycetes | I2H5M2 | I2H871 | I2H4Z2 |
| 35 | *Tetraodon nigroviridis* | Chordata | Q4RRE1* | H3DHR7 | H3C4U1* |
| 36 | *Tuber melanosporum* | Ascomycetes | D5GLY4 | D5GKM5 | D5G5D6 |
| 37 | *Saccharomyces cerevisiae* | Ascomycetes | P40368 | P14907 | P40477 |
| 38 | *Xenopus laevis* | Chordata | Q4KLQ6 | Q91349 | Q9PVZ2 |

*Fragment amino acid sequence available

### Figure S3

#### a) Nup88

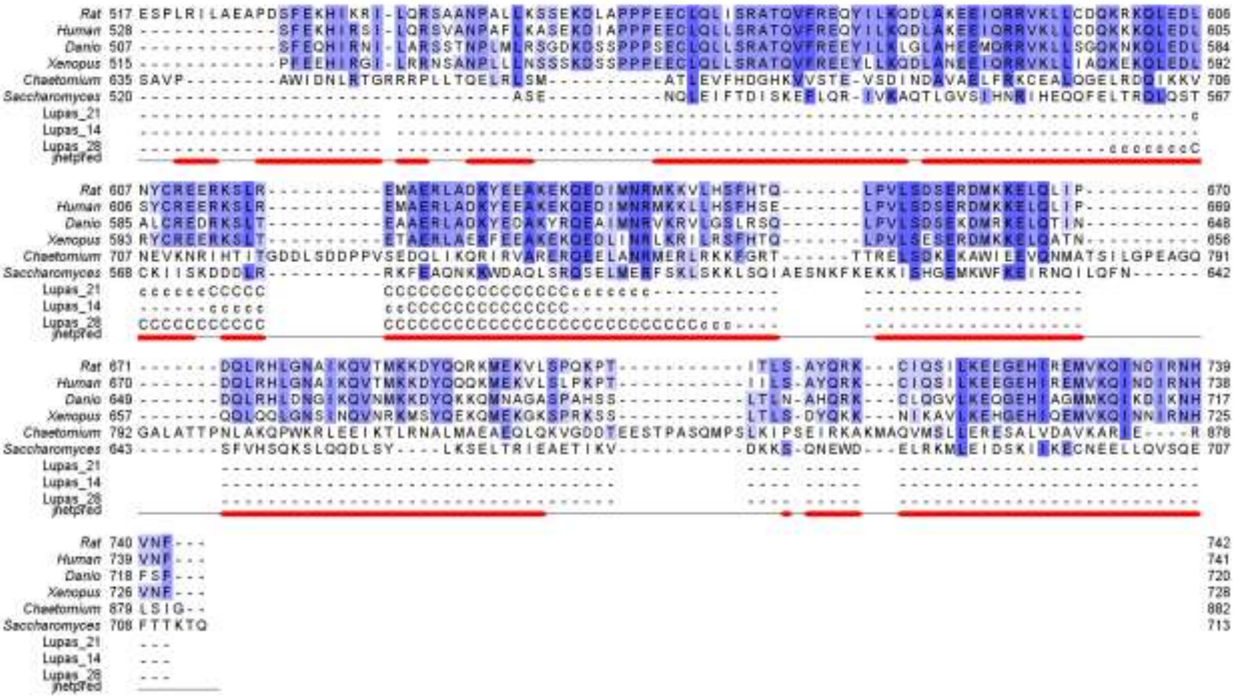

#### b) Nup62

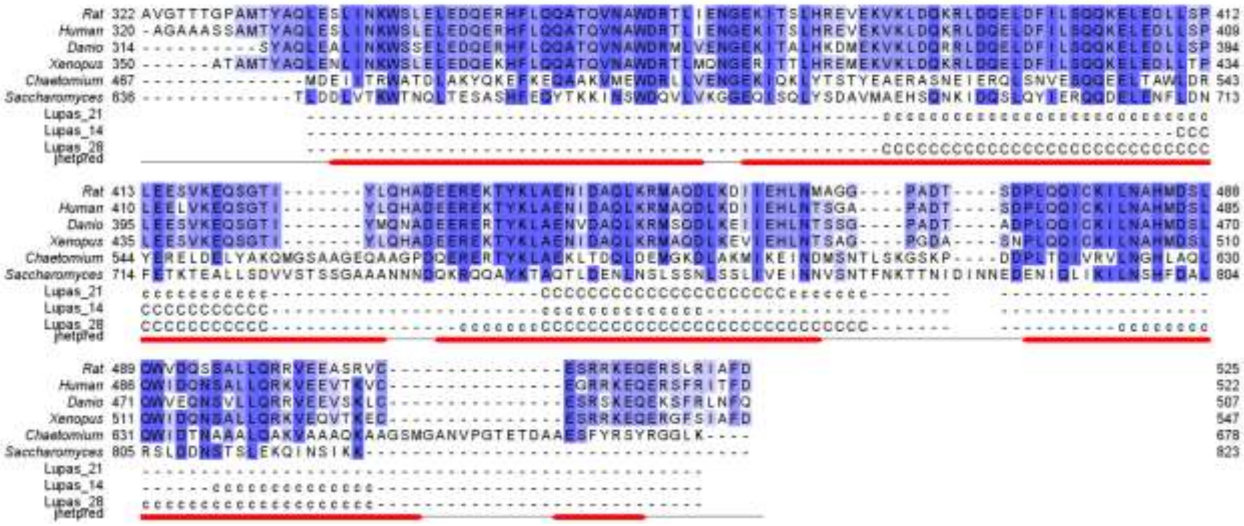

##### c) Nup214

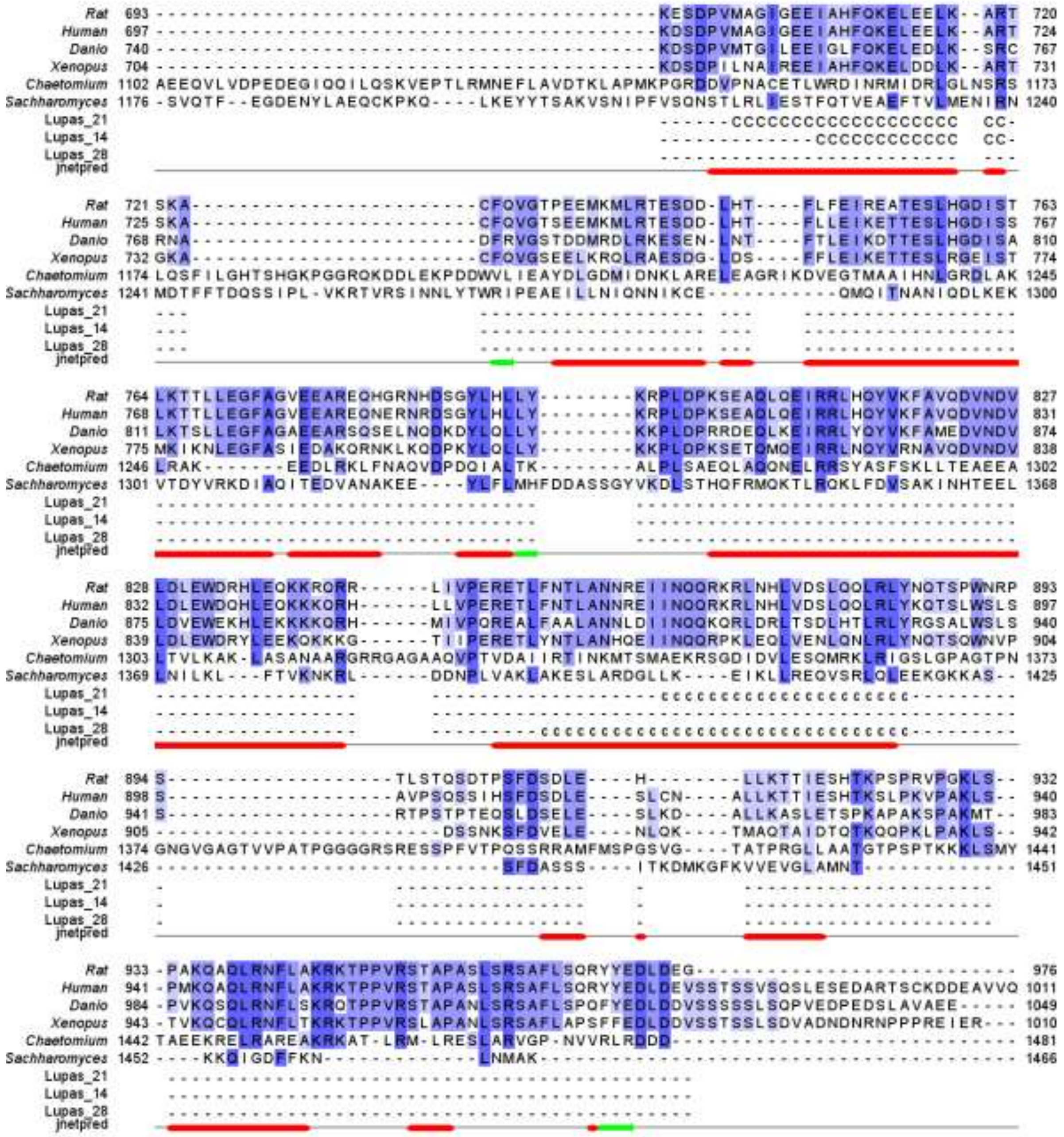


**Fig. S3: Structure guided alignment of (a) Nup88, (b) Nup62 and Nup214 protein sequences:** Multiple sequences aligned using PROMALS3D and viewed using Jalview. The cloned coiled-coil region is compared for six different species. Conserved secondary structure domains viewed with Jpred and coiled-coil motifs were predicted using Coils integrated in the work-flow of the Jalview.

**Table S2: Percent Identity Matrix (PIM) for coiled-coil regions of Nup88, Nup62 and Nup214 from different species:**

**a) Nup88/Nup82 coiled-coil:**

| | **1** | **2** | **3** | **4** | **5** | **6** |
|---|---|---|---|---|---|---|
| **1. *S. cerevisiae* Nup82** | 100.00 | 21.35 | 16.94 | 16.94 | 17.74 | 18.55 |
| **2. *C. thermophilum* Nup82** | 21.35 | 100.00 | 20.21 | 20.21 | 22.28 | 21.24 |
| **3. *D. rerio* Nup88** | 16.94 | 20.21 | 100.00 | 63.93 | 66.06 | 64.71 |
| **4. *X. laevis* Nup88** | 16.94 | 20.21 | 63.93 | 100.00 | 70.32 | 68.49 |
| **5. *R. rattus* Nup88** | 17.74 | 22.28 | 66.06 | 70.32 | 100.00 | 91.89 |
| **6. *H. sapiens* Nup88** | 18.55 | 21.24 | 64.71 | 68.49 | 91.89 | 100.00 |

**b) Nup62/Nsp1 coiled-coil:**

| | **1** | **2** | **3** | **4** | **5** | **6** |
|---|---|---|---|---|---|---|
| **1. *S. cerevisiae* Nsp1** | 100.00 | 32.63 | 31.28 | 29.05 | 28.49 | 29.61 |
| **2. *C. thermophilum* Nsp1** | 32.63 | 100.00 | 34.52 | 32.99 | 32.99 | 34.01 |
| **3. *D. rerio* Nup62** | 31.28 | 34.52 | 100.00 | 79.80 | 82.83 | 82.32 |
| **4. *X. laevis* Nup62** | 29.05 | 32.99 | 79.80 | 100.00 | 85.00 | 87.00 |
| **5. *R. rattus* Nup62** | 28.49 | 32.99 | 82.83 | 85.00 | 100.00 | 90.24 |
| **6. *H. sapiens* Nup62** | 29.61 | 34.01 | 82.32 | 87.00 | 90.24 | 100.00 |

**c) Nup214/Nup159 coiled-coil:**

| | **1** | **2** | **3** | **4** | **5** | **6** |
|---|---|---|---|---|---|---|
| **1. *S. cerevisiae* Nup159** | 100.00 | 16.03 | 15.15 | 13.85 | 14.72 | 14.91 |
| **2. *C. thermophilum* Nup159** | 16.03 | 100.00 | 19.79 | 20.49 | 17.33 | 17.60 |

| | | | | | | |
|---|---|---|---|---|---|---|
| **3. *D. rerio* Nup214** | 15.15 | 19.79 | 100.00 | 52.98 | 61.17 | 63.44 |
| **4. *X. laevis* Nup214** | 13.85 | 20.49 | 52.98 | 100.00 | 63.14 | 60.21 |
| **5. *R. rattus* Nup214** | 14.72 | 17.33 | 61.17 | 63.14 | 100.00 | 88.73 |
| **6. *H. sapiens* Nup214** | 14.91 | 17.60 | 63.44 | 60.21 | 88.73 | 100.00 |

### Figure S4

#### a) Protein pair Nup214 and Nup88

| Nup214 | Nup88 | Convolution Scores |
|---|---|---|
| 950-985 | 304-308 | 452 |
| 10-27 | 304-308 | 372 |
| 1006-1017 | 304-308 | 248 |
| 1091-1099 | 304-308 | 222 |
| 343-351 | 611-617 | 204 |
| 934-947 | 304-308 | 194 |
| 277-282 | 620-628 | 186 |
| 750-760 | 304-308 | 179 |
| 1091-1100 | 623-625 | 173 |
| 201-205 | 611-617 | 167 |

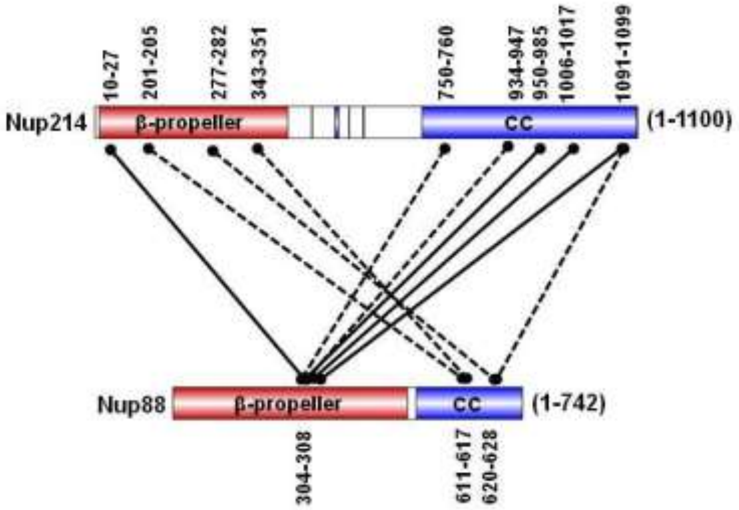

**b) Protein pair Nup214 and Nup62**

| Nup214 | Nup62 | Convolution Scores |
|---|---|---|
| 507-524 | 370-374 | 223 |
| 1007-1020 | 488-491 | 209 |
| 838-844 | 424-429 | 112 |
| 574-578 | 394-398 | 102 |
| 1069-1072 | 370-374 | 80 |
| 839-844 | 346-349 | 79 |
| 513-520 | 395-398 | 78 |
| 1007-1015 | 481-483 | 76 |
| 424-429 | 403-405 | 72 |
| 971-977 | 454-456 | 68 |

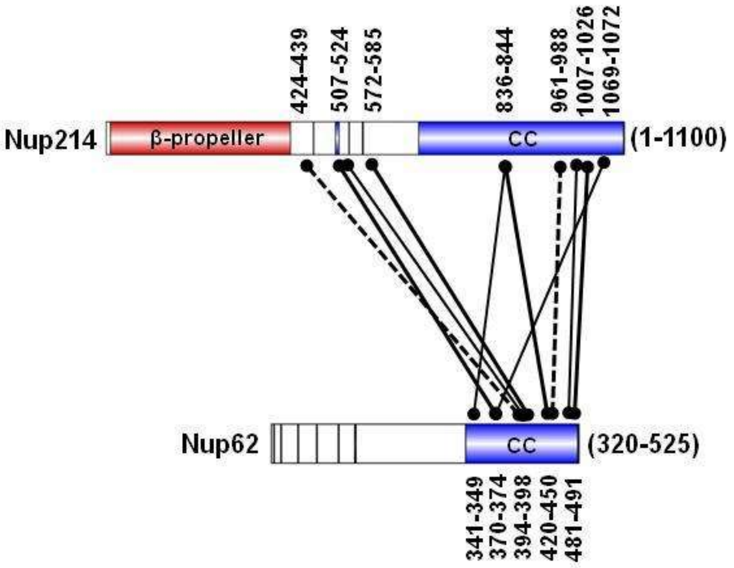

c) **Protein pair Nup62 and Nup88**

| Nup62 | Nup88 | Convolution Scores |
|---|---|---|
| 370-373 | 140-152 | 215 |
| 370-373 | 609-618 | 205 |
| 488-492 | 612-626 | 195 |
| 364-366 | 610-626 | 174 |
| 481-484 | 614-626 | 163 |
| 394-398 | 609-618 | 160 |
| 340-343 | 142-152 | 146 |
| 405-407 | 620-631 | 85 |
| 431-433 | 304-310 | 71 |
| 481-484 | 564-574 | 61 |

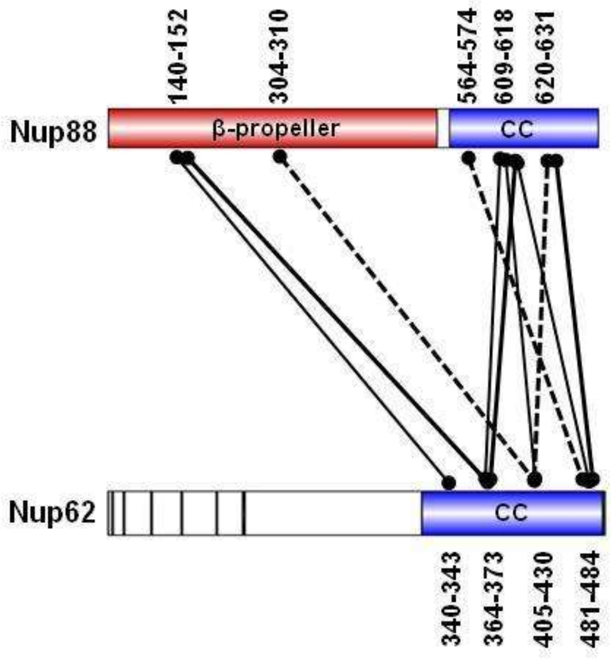

**Fig. S4: CoRNeA predictions for inter-domain interaction network of Nup88 complex** (related to Fig. 2a): CoRNeA tools were used to predict the interaction network between various regions of Nup88, Nup62 and Nup214. Piar-wise representation of the interaction network between Nups has been shown (a-c) and the corresponding tables represent the predicted convoluted score value. The scores marked in red color are considered as ***high*** and denoted by solid black lines (**━**); ***intermediate*** scores are shown in blue color and indicated by thin black lines (—) while ***low*** scores are highlighted in green color and marked by dashed lines (·····).

### Figure S5

#### a) In-gel trypsin-digested Nup88 complex

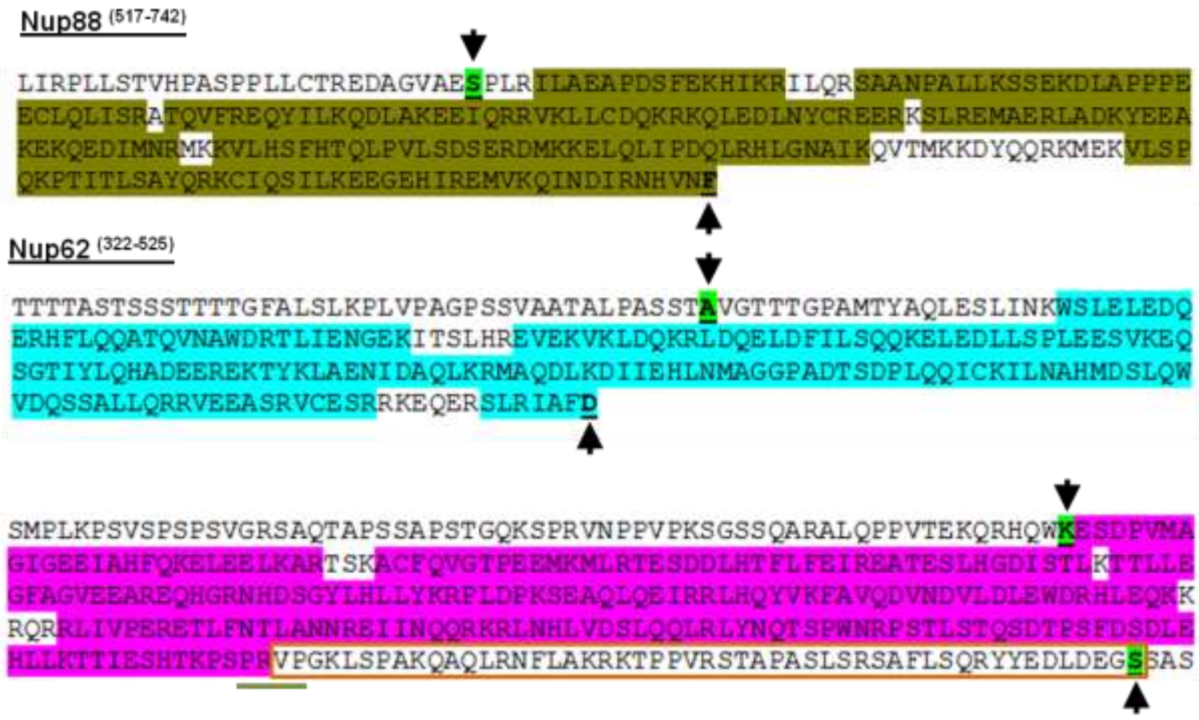


#### b) In-sol trypsin-digested Nup88 complex

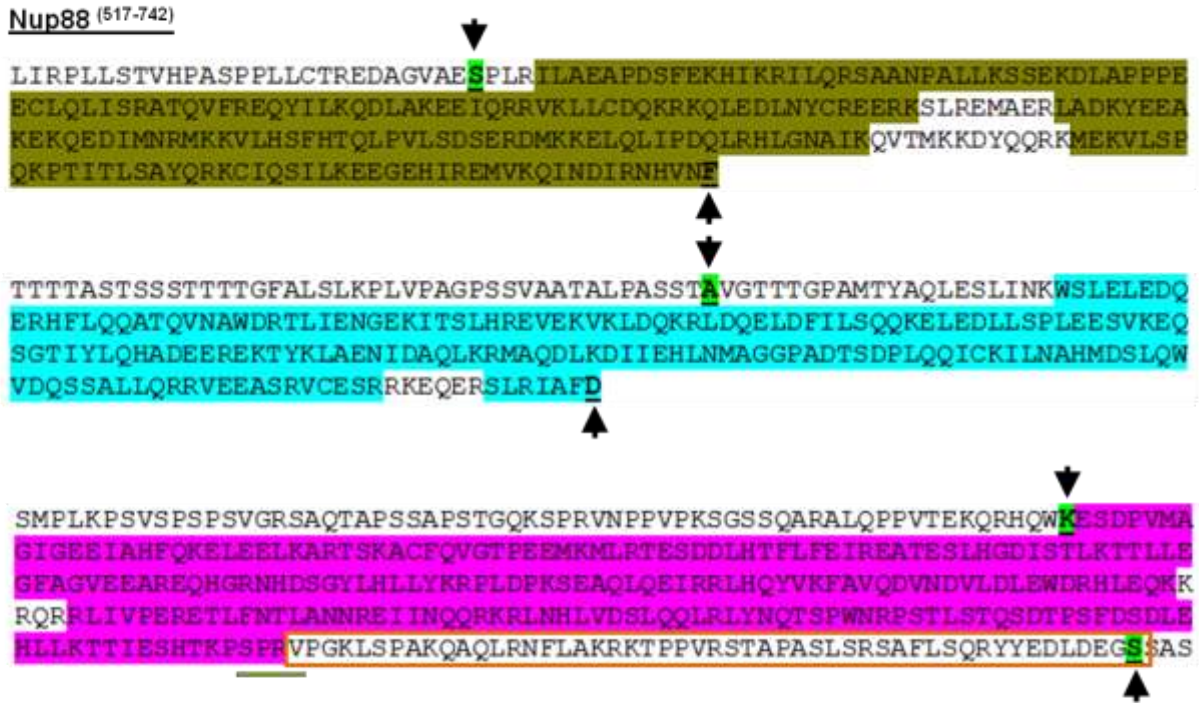

**Fig. S5: Mass spectrometry analysis**: The arrows mark the start and end residues of the cloned and expressed protein. The color highlighted regions show the overlapping fragments obtained as hits in Orbitrap MS analysis. The boxed region (red color) in Nup214 (in both In-gel and In-sol digestion) represents the region which has not been covered in Mass Spectrometry. This could be the region which was cleaved off during thrombin digestion and so the molecular weight of the digested protein is reduced by 6 kDa. The residues underlined (green) in the Nup214 sequence might serve as non-specific site for thrombin cleavage.

**<u>Table S3:</u> Orbitrap LC-MS search result summary for trypsin-digested Nup88 complex**

**(*Top hits with maximum scores are highlighted in red*)**

| In-gel trypsin digestion | | | | | |
|---|---|---|---|---|---|
| Protein FDR Confidence | Description | Coverage [%] | Peptides | Unique Peptides | Score Mascot |
| High | Nuclear pore complex protein Nup88 OS=Rattus norvegicus OX=10116 GN=Nup88 PE=1 SV=1 | 27 | 48 | 48 | 11251 |
| High | Nucleoporin 214 (Fragment) OS=Rattus norvegicus OX=10116 GN=Nup214 PE=1 SV=2 | 13 | 29 | 29 | 9122 |
| High | Nuclear pore glycoprotein p62 OS=Rattus norvegicus OX=10116 GN=Nup62 PE=1 SV=1 | 33 | 24 | 24 | 8082 |
| High | Keratin, type II cytoskeletal 5 OS=Rattus norvegicus OX=10116 GN=Krt5 PE=1 SV=1 | 12 | 10 | 6 | 232 |
| High | Keratin, type I cytoskeletal 10 OS=Rattus norvegicus OX=10116 GN=Krt10 PE=1 SV=1 | 10 | 6 | 4 | 250 |
| High | Cytokeratin-1 OS=Rattus norvegicus OX=10116 GN=Krt1 PE=1 SV=1 | 5 | 5 | 3 | 261 |
| High | Keratin, type II cytoskeletal 2 epidermal OS=Rattus norvegicus OX=10116 GN=Krt2 PE=3 SV=1 | 6 | 4 | 3 | 151 |

| In-solution trypsin digestion | | | | | |
|---|---|---|---|---|---|
| Protein FDR Confidence | Description | Coverage [%] | Peptides | Unique Peptides | Score Mascot |
| High | Nuclear pore complex protein Nup88 OS=Rattus norvegicus OX=10116 GN=Nup88 PE=1 SV=1 | 27 | 56 | 56 | 12931 |
| High | Nucleoporin 214 (Fragment) OS=Rattus norvegicus OX=10116 GN=Nup214 PE=1 SV=2 | 13 | 38 | 38 | 12081 |
| High | Nuclear pore glycoprotein p62 OS=Rattus norvegicus OX=10116 GN=Nup62 PE=1 SV=1 | 34 | 28 | 28 | 9064 |
| High | Keratin, type II cytoskeletal 5 OS=Rattus norvegicus OX=10116 GN=Krt5 PE=1 SV=1 | 14 | 11 | 5 | 365 |
| High | Cytokeratin-1 OS=Rattus norvegicus OX=10116 GN=Krt1 PE=1 SV=1 | 7 | 6 | 4 | 498 |
| High | Keratin, type II cytoskeletal 2 epidermal OS=Rattus norvegicus OX=10116 GN=Krt2 PE=3 SV=1 | 8 | 6 | 3 | 351 |
| High | Keratin, type II cytoskeletal 6A OS=Rattus norvegicus OX=10116 GN=Krt6a PE=1 SV=1 | 11 | 7 | 2 | 288 |

| | | | | | |
|---|---|---|---|---|---|
| High | Keratin, type I cytoskeletal 15 OS=Rattus norvegicus OX=10116 GN=Krt15 PE=1 SV=1 | 6 | 6 | 5 | 244 |
| High | Keratin, type I cytoskeletal 10 OS=Rattus norvegicus OX=10116 GN=Krt10 PE=3 SV=1 | 12 | 6 | 5 | 211 |
| High | Heat shock-related 70 kDa protein 2 OS=Rattus norvegicus OX=10116 GN=Hspa2 PE=1 SV=1 | 7 | 3 | 2 | 203 |
| High | Desmoplakin OS=Rattus norvegicus OX=10116 GN=Dsp PE=1 SV=1 | 2 | 5 | 5 | 162 |
| High | Keratin 16 OS=Rattus norvegicus OX=10116 GN=Krt16 PE=1 SV=1 | 15 | 5 | 4 | 139 |

**Figure S6**

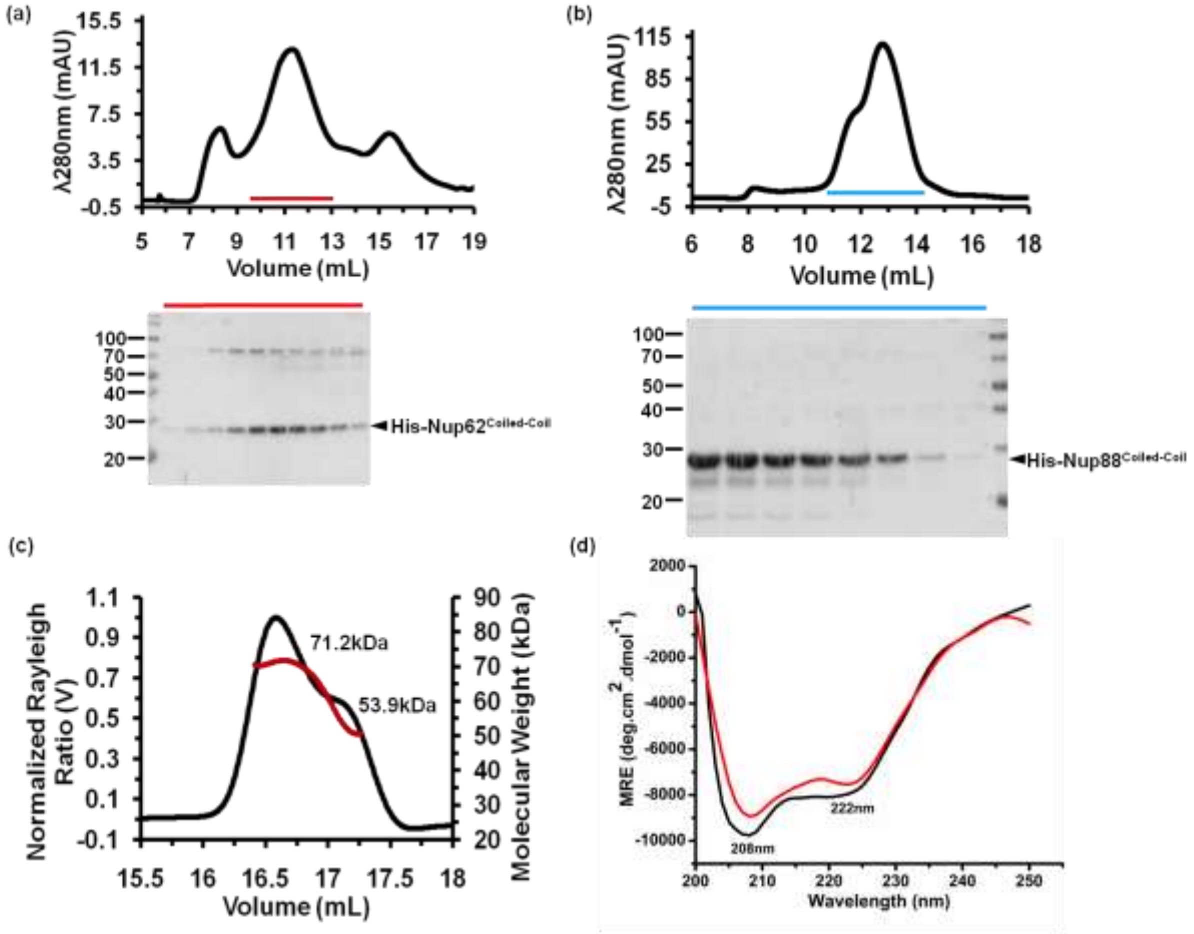


**Fig. S6: Purification of Nup62 and Nup88 alone:** (a) SEC profile and SDS-PAGE for the Nup62 coiled-coil region (a), and Nup88 coiled-coil (b). The fractions marked in red and blue color were loaded on 12% reducing PAGE which shows bands of Nup62 and Nup88, respectively. (c) SEC-MALS of Nup88 indicating a dynamic equilibrium between trimer and dimer state of the protein. The molar mass ranges from 71kDa to 53kDa as marked in red color (also see table S5). (d) Far UV CD spectra for both Nup62 (black) and Nup88 (red) which suggest a typical α-helical profile with two signature minima at 208 nm and 212 nm respectively.

**<u>Table S4:</u> Theoretical and estimated molar mass calculated by SEC-MALS for Nup88 trimeric (Nup88•Nup214•Nup62) complex, Nup88 dimeric (Nup88•Nup62) complex and Nup88 alone**

| | **Nup88•Nup214•Nup62** | **Nup88•Nup62** | **Nup88** |
|---|---|---|---|
| **Protein Concentration (mg/mL)** | 1.4 | 0.9 | 0.8 |
| **Calculated Molecular Weight** | 1.4 mg/ml: 69.3 (±0.3%) (0.4 mg/ml: 68.3) | 71.1 (±0.7%) | 65.3 (±0.6%) (Peak1: 71.2 Peak2: 53.9) |
| **Theoretical Molecular Weight (kDa)** | 77.5 | 52.8 | 29.1 |
| **Possible Stoichiometry (ratio)** | 1:1:1 | 1:2 | --- |

**Figure S7**

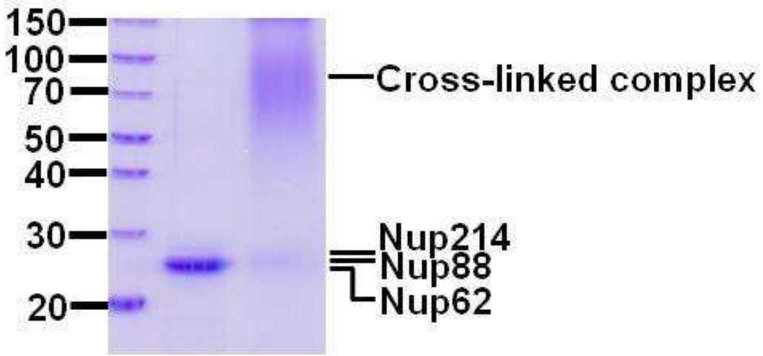


**Fig. S7:** Chemical cross linking (0.1% Glutaraldehyde, 15 min) of the Nup88•Nup214•Nup62 heterotrimeric complex shows smeary band near 70-90 kDa supporting the SEC-MALS data. The absence of a sharp band could be due the dynamic behaviour of the complex. Lane (-) is the control with no cross linker showing the complex (without tag). Lane (+) marks the cross linked complex on 12% SDS PAGE.

**Figure S8**

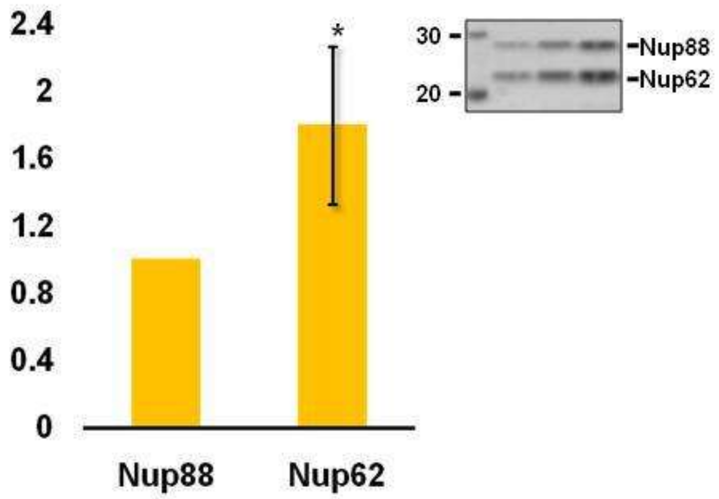


**Fig. S8:** Densitometric analysis of the purified Nup88•Nup62 protein complex bands show almost double band intensity for Nup62 than Nup88. Protein bands are shown in the inset.

#### Figure S9

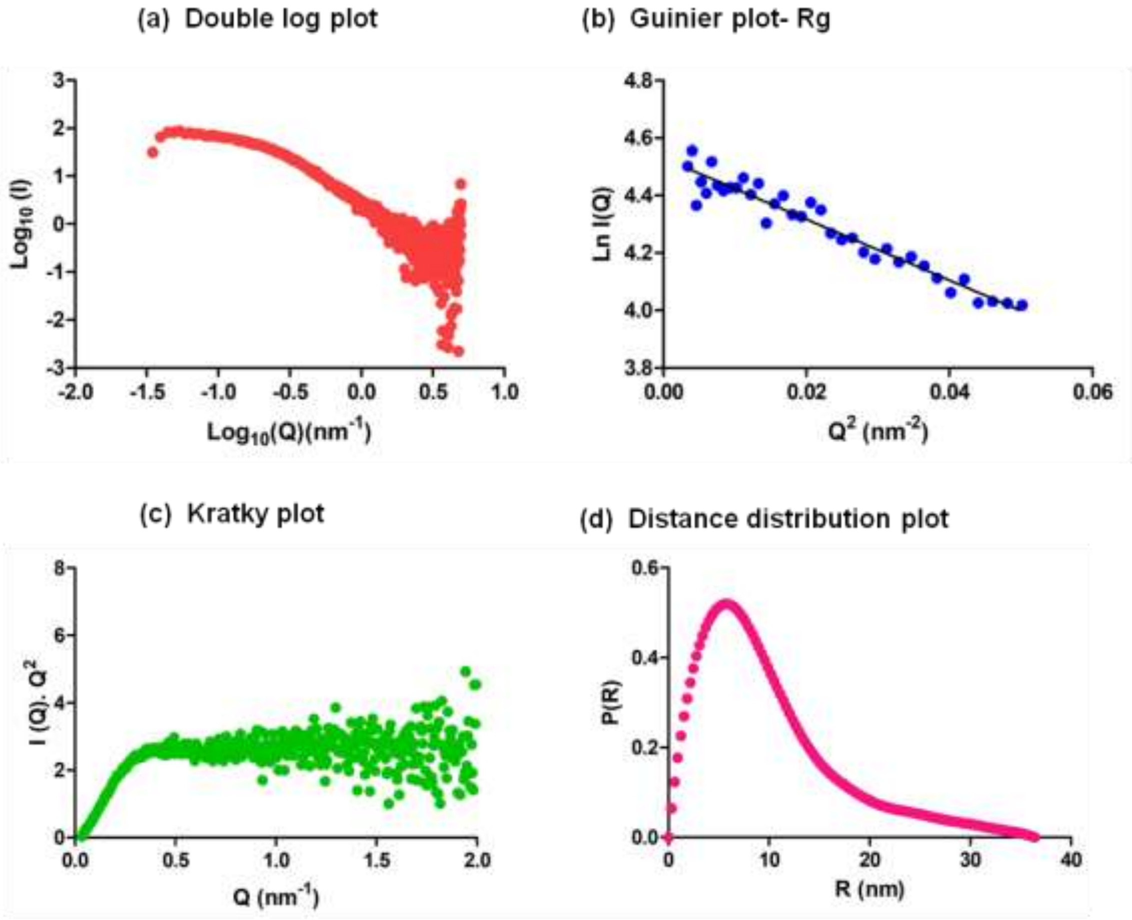


**Fig. S9: Small Angle X-ray Scattering (SAXS) analysis of Nup88$^{517\text{-}742}$•Nup62$^{322\text{-}525}$** (related to Fig. 6)**:** (a) Double log plot Log10 Q vs. Log10 I (Q) of the complex. (b) Guinier plot analysis: Ln I (Q) vs $Q^2$ ($nm^{-2}$). (c) Kratky analysis by plotting I (Q)•$Q^2$vs (Q) for the complex. (d) Pair-wise distribution analysis P(r) the complex.

#### Table S5

**a: Ratio of the molar ellipticity at 222 nm and 208 nm and % helicity of the complexes:**

| **Complexes** | $\theta_{222}/\theta_{208}$ | **% Helicity** |
|---|---|---|
| Nup62•Nup88 dimeric complex | 0.97 | 38.4 |
| Nup62•Nup88•Nup214 trimeric complex | 1.00 | 47.5 |

**b: Percent helicity of the complexes at different temperatures:**

| **Temperature (°C)** | **% Helicity** | |
|---|---|---|
| | **Dimeric complex** | **Trimeric complex** |
| **25** | 38.4 | 47.5 |
| **30** | 37.8 | 47.5 |
| **40** | 36.2 | 44.9 |
| **50** | 33.0 | 44.9 |
| **60** | 28.0 | 37.2 |

<u>**Figure S10**</u>

(a)

| Parameter | |
|---|---|
| Microscope | JEM-2200FS |
| Magnification | 20000 |
| Voltage (kV) | 200 |
| Detector | Gatan camera (K2 summit direct detection camera) |
| Sample staining | 2% Uranyl acetate |
| Pixel Size (Å) | 1.9 |
| No. of Micrographs | 55 |
| No. of particles used for reconstruction | 4737 |
| Map resolution (Å)<br>@FSC criteria | 19.27<br>0.143 |

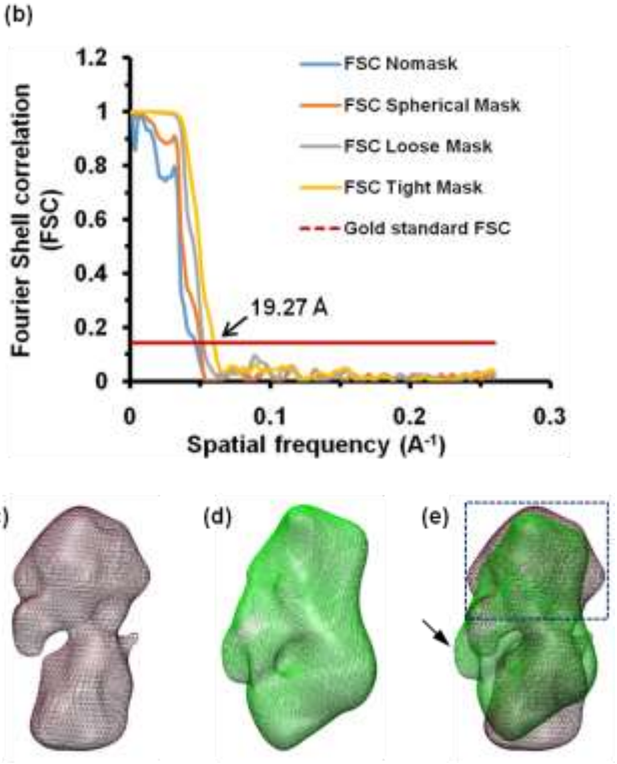


**Fig. S10: EM analysis of rNup88 (Nup88$^{517-742}$•Nup62$^{322-525}$•Nup214$^{693-976}$) complex** (related to Fig. 8): (a) Summary of the EM data collection and analysis. (b) FSC curve for the refined 3D map of the Nup88 complex, showing the resolution of EM map at 19.27 Å, indicated by arrow. Superposition of CTC•Nup93$^{1-150}$ complex over Nup88 complex: (c) 3D density map of Nup88 complex (shown in grey color), (d) 3D density map of CTC•Nup93 complex (shown in green color), (e) superimposed 3D density map of Nup88 complex (grey) over the CTC•Nup93 complex (green). Black arrow indicates the extra density for Nup93$^{1-150}$ while blue box shows the overlapping head region of both the complexes.

**Figure S11**

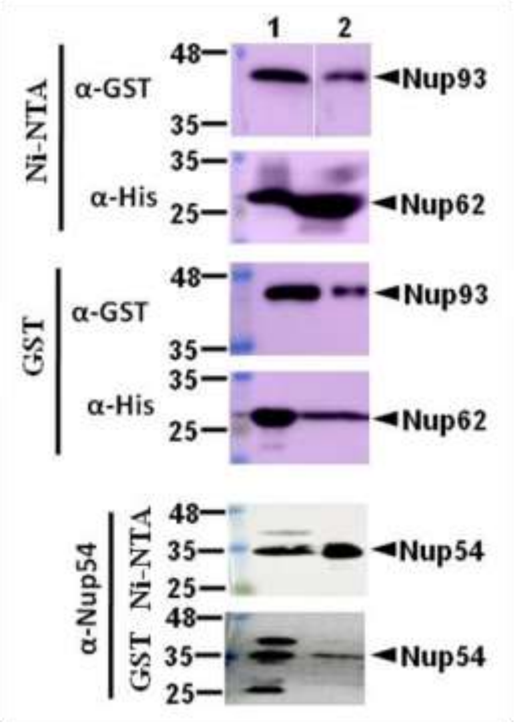


**Fig. S11:** (related to figure 9): Western analysis of reconstituted Nup62•Nup54•Nup93 complex which shows that the N-terminal domain (1-150) of Nup93 interacts with Nup62 and Nup54. The co-eluted fractions were probed with anti-His, anti-GST and anti-Nup54 antibodies. Lane 1, 2 represent input and eluted fraction respectively.

**<u>Table S6</u>: Details of expression plasmids and constructs used in the study**

| Protein | Expression vector | Cloning sites | Primers used for cloning (5’-3’) |
|---|---|---|---|
| *Rat*Nup88 (517-742) and *Rat*Nup62 (322-525) | pRSF-Duet1 | *SalI* | F:CAGGTCGACAGCAGCGGCCTGGTGCCGCGCGGCAGCGAGTCTCCACTGCGCATCCTGG |
| | | *HindIII* | R:CGCAAGCTTTTATCAGAAGTTGACATGATTTCGG |
| | | *NdeI* | F:CCTACATATGGGGACTACGACGGGTCCAGCAATG |
| | | *XhoI* | R:AGACTCGAGTTACTAGTCAAAGGCAATGCGCAGG |
| *Rat*Nup214 (693-976) | pGEX-4T1 | *BamHI* | F:CGTGGATCCATGAAAGAGTCAGACCCTGTGATGG |
| | | *XhoI* | R:CCGCTCGAGTTATCAGCCTTCATCTAAGTCTTCATAG |
| *Rat*Nup214 (1-407) | pGEX-4T1 | *EcoRI* | F: CGTGAATTCATGGGAGACGAGATGGATGC |
| | | *SalI* | R:TGCG*GTCGAC*TCAATTTTGATTGATCATATAAAATGGAC |
| *Rat*Nup88 (517-742) | pET-28a | *EcoRI* | F:CCGGAATTCGAGTCTCCACTGCGCATCCTGG |
| | | *HindIII* | R:CGCAAGCTTTTATCAGAAGTTGACATGATTTCGG |
| *Rat*Nup88 (59-498) | pET-28a | *EcoRI* | F: CCGGAATTCACGAGAAACCTGGTCTTCGGC |
| | | *HindIII* | R:CGCAAGCTTTTATCATGTACTTAATAAAGGCCTTATGAGAC |
| *Rat*Nup62 | pGEX-4T1 | *BamHI* | F:CGCGGATCCGGGACTACGACGGGTCCAGCAATG |

| | | | |
|---|---|---|---|
| (322-525) | | *XhoI* | R:AGACTCGAGTTACTAGTCAAAGGCAATGCG CAGG |
| Human Nup54 and Human Nup93 | pET Duet1 | *NcoI* | CATGCCATGGGCATGCCCAGTAATAAAGATG AAGATGGG |
| | | *HindIII* | CCCAAGCTTTCAACTAAAGACACCACCTCTGA TGTG |
| | | *NdeI* | GGAATTCCATATGTCCCCTATACTAGGTTATT GG |
| | | *XhoI* | CCGCTCGAGTTAATTCATGAGGACCTCCAT |
